# Gradient Multinozzle 3D Printing

**DOI:** 10.64898/2026.03.21.712762

**Authors:** Luca Rosalia, Soham Sinha, Jonathan D. Weiss, Senming Hsia, Fredrik S. Solberg, Amit Sharir, Masufumi Shibata, Jianyi Du, Katelyn Mosle, Dominic Rütsche, Zi Chang Rao, Tony Tam, Trenton Rankin, Qiuling Wang, Christian M. Williams, John H. Klich, Alexander K. Reed, Eric A. Appel, Michael Ma, Mark A. Skylar-Scott

**Author notes:** These authors contributed equally to this work (Luca Rosalia, Soham Sinha, Jonathan D. Weiss).

## Abstract

Direct ink writing is compatible with an expansive materials palette. While enabling diverse applications, this materials versatility brings significant bottlenecks in ink formulation, often requiring the mixing, printing, and testing of dozens to hundreds of ink compositions over the course of a project. To accelerate ink-space exploration, we introduce gradient embedded multinozzle (GEM) printheads that combine the high-throughput parallelized printing of multinozzles with combinatorial ink mixing. These printheads allow simultaneous mixing of two-, three-, and four-input inks which are distributed to printer nozzles to create complex 3D structures with graded compositions of inks. Using a two-way GEM printhead, we validate cell compatibility by printing scaffolds containing various concentrations of fibroblasts and observing non-linear compaction behaviours. We next test a three-way GEM multinozzle to print ten compositions of di- and multi-functionalized poly(ethylene-glycol) diacrylate hydrogel tri-leaflet valves, optimizing for stiffness, swelling ratio, and toughness. Our GEM multinozzles are compatible with open-source printers and either pressure- or volume-driven extrusion systems and promise to accelerate iterative ink design and testing.

## 1 Introduction

Direct ink writing (DIW) is a highly versatile method of 3D printing with wide ranging materials from printed composites^1–3^, metals^4–7^, ceramics^8–10^, electronics^5,11,12^ and biological^13–15^ materials. DIW depends upon the formulation of viscoelastic inks that are both printable and impart functional properties^16^. Ink optimization is the most time-intensive step in developing new DIW processes due to the large combinatorial space of formulation parameters. For hydrogels and bioinks, this typically involves tuning polymer composition, crosslinking chemistry, rheological properties, and biological components such as extracellular matrices, cell types, and growth factors. Rather than exhaustive multidimensional searches, this process is often guided by empirical intuition. Furthermore, formulating viscoelastic, yield-stress inks for DIW requires laborious handling, including mixing, centrifugation, syringe loading, and degassing, which can become prohibitively time consuming and result in material loss during transfer steps. This is especially limiting for inks that have short pot lives or those containing living cells, where extended processing times can compromise reproducibility and cell viability.

Eliminating manual formulation steps and parallelizing construct fabrication can substantially accelerate 3D-printed assays. Existing multinozzle printheads enable parallel printing of single^17^ or multiple^18,19^ materials, but all nozzles print identical ink compositions at any given time, yielding arrays of compositionally equivalent constructs. Mixing printheads can generate different ratios of, generally, two input materials^20,21^, but are limited to printing single constructs serially. Thus, current printhead technologies are not well-suited for large, parallel screens of graded constructs, particularly when inks have short pot lives^22^. An ideal printhead for formulation screening combines these capabilities by enabling parallelized mixing and printing of distinct ink compositions across an array of nozzles. By printing many combinations of material mixtures simultaneously, an optimal formulation that exhibits desired mechanical or biological properties may be quickly identified.

Here, we introduce gradient embedded multinozzle (GEM) 3D printheads that mix two, three, and four separate inks into distinct formulations and print them in parallel through an array of nozzles. GEM printheads employ bifurcating, trifurcating, or quadrifurcating channels interspersed with 3D baker’s map mixers^23^. Unlike conventional gradient microfluidic systems, which recombine mixed streams into a single outlet^24–26^, each mixed output is routed to an independent parallel printer nozzle.

The two-way, three-way, and four-way GEM printheads have eight, ten, and sixteen output nozzles, respectively. We apply the two-way printhead to create tissue scaffolds with different cellular densities and the three-way printhead to create tri-leaflet valves with different polymer blends. To facilitate distribution, the printheads are compatible with the open-source and low-cost Printess 3D printer^19^ and are distributed as fully editable, variable-driven design files with accompanying assembly and usage instructions (**Zenodo 10.5281/zenodo.17991829, Supplementary Fig. 1**).

## 2 Results and Discussion

### 2.1 Gradient Embedded Multinozzle Printheads

GEM 3D printheads are designed to perform embedded 3D printing of an array of identical 3D geometries, each composed of graded compositions of input materials (**Fig. 1a–c, Supplementary Video 1**). The printheads, manufactured using stereolithography, contain complex internal geometries that combine, mix, and redistribute input materials toward an array of output nozzles. We demonstrate two-way, three-way, and four-way mixing printheads with eight, ten, and sixteen nozzles, respectively (**Fig. 1d–f**). Graded arrays of colored rings can be printed by loading the printheads’ inlet syringes with Carbopol (0.3 wt% ETD 2020), a yield-stress and shear-thinning ink, containing differing fluorescent pigments (**Fig. 1g**). GEM printheads are compatible with direct ink writing methods that employ either pressure-based (**Fig. 1a–c**) or volume-based (**Fig. 1g**) extrusion. We additionally provide a geared extruder design that facilitates adoption on the low-cost Printess 3D printer (**Supplementary Fig. 2**).

**Fig. 1:**
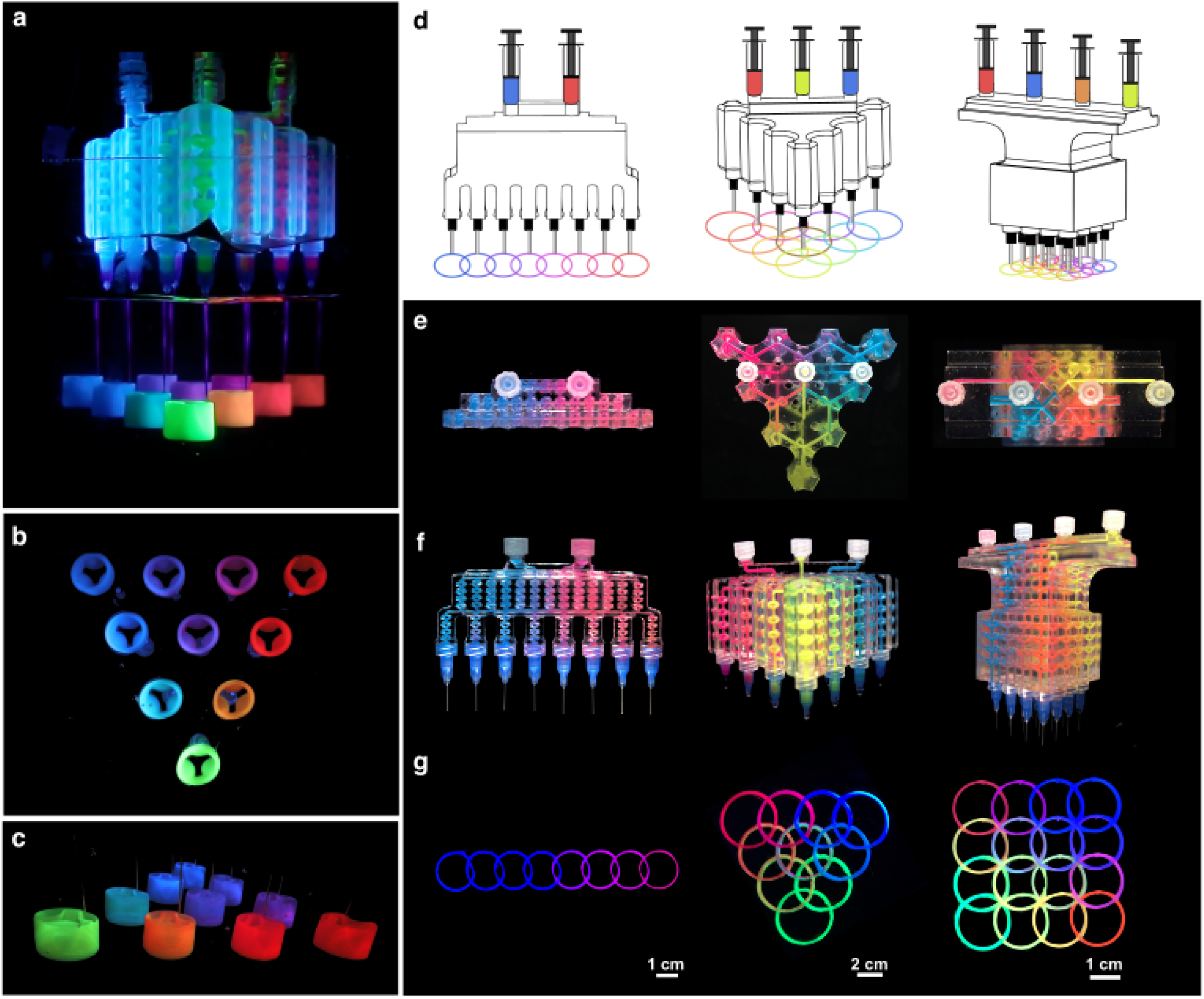
Two-, three-, and four-way gradient embedded multinozzle printheads. **a–c**, Photography of trileaflet valve printing using the three-way printhead shown in front view (**a**), bottom view (**b**), and isometric view (**c**). **d**, Schematics of two-, three-, and four-way gradient embedded multinozzle printheads. **e–f**, Photography of two-, three-, and four-way gradient embedded multinozzle printheads shown in top view (**e**) and front or isometric view (**f**). **g**, Array of two-dimensional rings printed using the two-, three-, and four-way gradient embedded multinozzle printheads.

### 2.2 Characterization of Gradient Embedded Multinozzle Printheads

Active and passive mixers have been developed to homogenize viscoelastic inks via active^27^ or passive^28^ mechanisms. While active mixers enable efficient mixing in junctions with small volumes, their associated electronics become cumbersome when scaling toward printheads containing dozens of junctions. Chaotic advection mixers provide an effective passive alternative for highly viscous, low-Reynolds-number materials, including polymeric and hydrogel inks used for 3D printing^29,30^.

GEM printheads employ passive mixers based on a baker’s map transformation geometry^31,32^, which repeatedly split and recombine input flows in orthogonal planes (**Fig. 2a–b, Supplementary Fig. 1**). This mechanism is well suited for mixing yield-stress Herschel–Bulkley fluids and produces progressively more uniform mixtures as additional mixing units are placed in succession (**Fig. 2c**). By approximately 6– 8 mixers, Carbopol (0.3 wt% ETD 2020) ink was mixed at the microscale, as confirmed by a homogeneous distribution of fluorescent microbeads at the outlet (**Fig. 2d, Supplementary Fig. 3**).

**Fig. 2:**
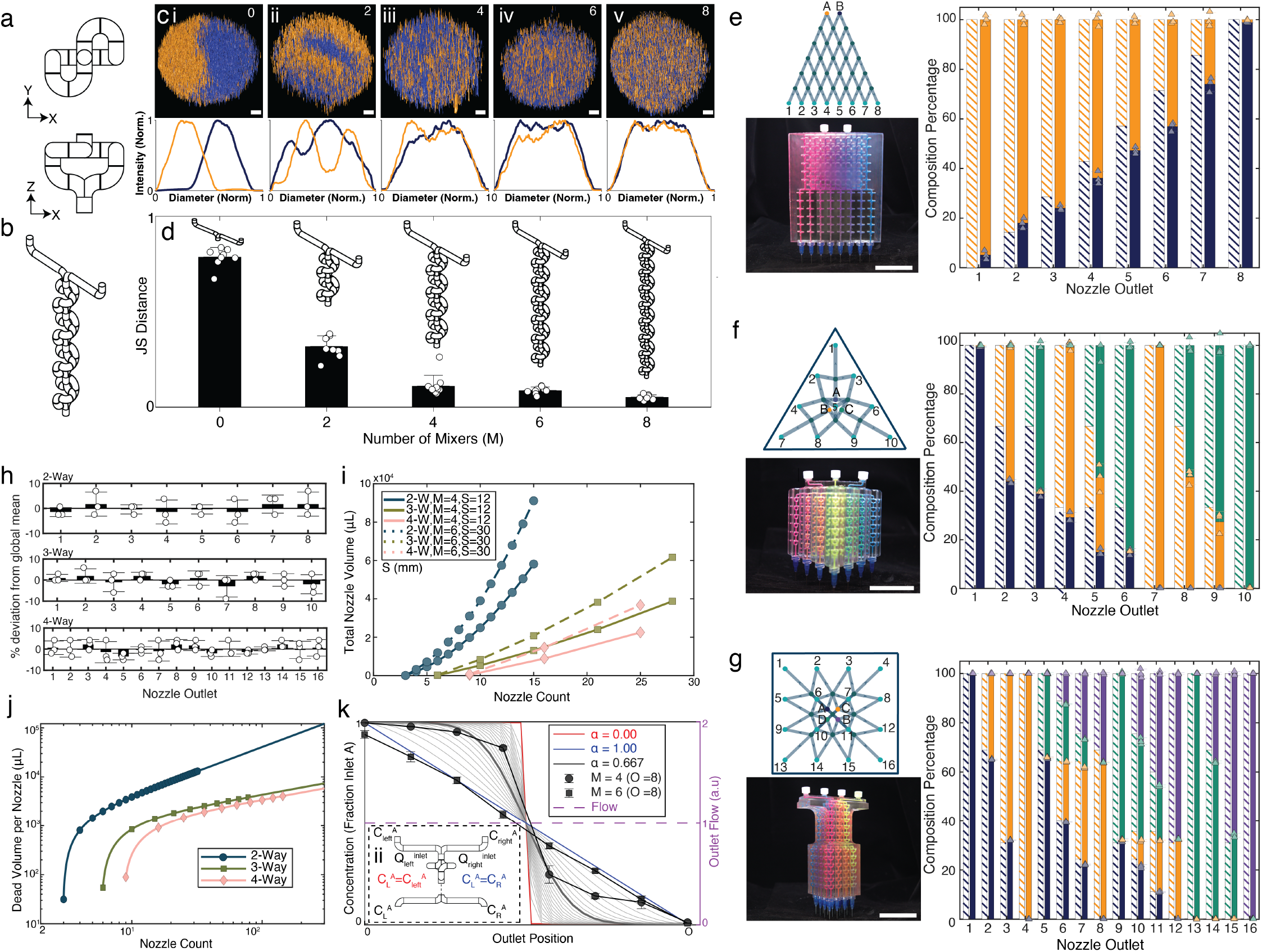
Characterization of two-, three-, and four-way gradient embedded multinozzle printheads. **a**, Planar views of a single mixing element in the XY and XZ planes. **b**, Isometric view of a four-mixer stack (*M* = 4) with two converging input streams. **c** (**i–v**), Confocal cross-sections of filaments printed with increasing numbers of mixers (0–8), with normalized diametrical fluorescence intensity profiles shown below (scale bar, 50 *µ*m). **d**, Jensen–Shannon distance between bead distributions as a function of mixer count (*n* = 9). **e–g**, Directed acyclic graph representations and experimental implementations of two-way (**e**), three-way (**f**), and four-way (**g**) GEM printheads (*M* = 6 variants shown; scale bars, 5 cm). For each architecture, predicted outlet compositions are shown as striped stacked bars and experimentally measured compositions as solid bars (*n* = 3). **h**, Flow-rate uniformity across nozzle outlets in two-, three-, and four-way printheads (*n* = 3). **i**, Total internal volume as a function of nozzle count showing different mixer numbers (*M* = 4, 6) and nozzle spacing (*S* = 12, 30 mm); markers indicate physically realizable designs. **j**, Dead volume per nozzle as a function of nozzle count for each printhead architecture (log–log scale, *S* = 12 mm, *M* = 6); markers indicate physically realizable designs. **k**, Incomplete mixing model for the two-way printhead showing outlet concentration profiles for varying mixing parameter *α* ∈ [0, 1]. Experimental data for *M* = 4 and *M* = 6 (eight outlets) are overlaid (*n* = 3). Extremes of *α* = 0 (no mixing) and *α* = 1 (perfect mixing) are highlighted; (**ii**) mass-balance model at a single mixing node.

Within a GEM printhead, input materials are combined at parent nodes, mixed via baker’s map transformations, and redistributed to downstream child nodes. The two-way printhead adopts a microfluidic Christmas-tree architecture commonly used for gradient generators^24,33^, in which the number of output nozzles grows linearly with the number of mixing layers and produces defined binary mixture ratios at the outlets (**Fig. 2e**).

The three-way printhead adopts a tetrahedral 3D architecture^34^, where nodes are positioned on a regular triangular lattice to generate defined ternary compositions (**Fig. 2f**). The four-way printhead adopts a square pyramidal architecture, with nodes arranged on a regular square lattice to generate defined quaternary compositions (**Fig. 2g**). When printing materials with equal rheology, outlet flows varied by less than approximately 10% across all three families (**Fig. 2h**), ensuring similar printed volumes.

To predict theoretical ink compositions at each outlet, we assume (i) equal rheologies for each inlet ink, (ii) symmetric total inflow across parent inlets, (iii) symmetric total outflow across outlets, (iv) mass conservation at all internal nodes, and (v) symmetric total outflow across nodes within the same level (**Supplementary Fig. 4**). Propagating inlet concentrations downstream using flux averaging yields an advective mixing matrix that predicts outlet composition fractions. Predicted concentrations closely matched experimentally measured values (**Fig. 2e–g**), obtained via confocal microscopy and spectral imaging of fluorescent microsphere distributions (**Supplementary Figs. 5, 6**).

The total dead volume of a GEM multinozzle is an important design consideration, particularly for expensive inks. Increasing nozzle spacing enables fabrication of larger constructs without intersection, while increasing the number of mixers improves mixing uniformity; however, both design choices increase dead volume (**Fig. 2i**). We therefore define a material efficiency metric as the total dead volume per nozzle, with lower values indicating more efficient material use. Material efficiency is substantially higher in three- and four-way designs because their nozzle count grows quadratically with the number of network layers, as opposed to linearly in two-way designs (**Fig. 2j**, Supplementary Text).

Incomplete mixing at each node can bias downstream redistribution, leading to unequal concentrations between child channels. To model this effect, we define a global mixing parameter *α*, representing the degree of mixing achieved at each two-way node. In this framework, *α* = 1 corresponds to perfect mixing and produces a linear gradient (**Fig. 2k**, blue line), while *α* = 0 corresponds to no mixing and yields a step function (**Fig. 2k**, red line). Intermediate values yield sigmoidal gradients of varying steepness (**Fig. 2k**, gray lines). Conceptually, *α* is analogous to fractional mixing coefficients used in universal gradient generators^35^ and is applied identically at each node, such that repeated application produces a global measure of effective mixing.

Consistent with this model, two-way printheads with *M* = 6 mixing elements exhibit near-linear outlet gradients (**Fig. 2k**, square datapoints), indicating near-complete mixing at each node, whereas printheads with fewer mixing elements (*M* = 4) exhibit sigmoidal gradients (**Fig. 2k**, circular datapoints). Outlet concentration distributions were independent of rheology, with near-identical gradients observed for Newtonian (glycerol) and Herschel–Bulkley (Carbopol) inks (**Supplementary Fig. 7**).

### 2.3 GEM 3D Printing of Tissue Constructs with Different Cellular Composition

In 3D bioprinting, the large combinatorial space of cell densities, cell types, extracellular matrix components, and rheological modifiers complicates systematic optimization of bioink formulations. This challenge is compounded by nonlinear and time-dependent cellular behavior in 3D scaffolds, which can produce sharp biological thresholds. For example, fibroblasts seeded in 3D collagen scaffolds exhibit a critical cell density at which cell–cell sensing triggers spreading, migration, and scaffold contraction^36^.

To validate GEM printheads for parallelized bioink formulation and exploration of such nonlinear regimes, we bioprinted fibrin scaffolds containing graded concentrations of human neonatal dermal fibroblasts (HNDFs). Two inks containing fibrinogen and gelatin, with either no cells or 1 million cells/mL, were prepared (**Fig. 3a**). Both inks exhibited similar consistency and shear-thinning behavior (**Fig. 3b, Supplementary Fig. 8a**). These inks were connected to a two-way GEM printhead and printed into a gelatin microgel support bath containing thrombin using Freeform Reversible Embedding of Suspended Hydrogels (FRESH) 3D printing^14,37^.

**Fig. 3:**
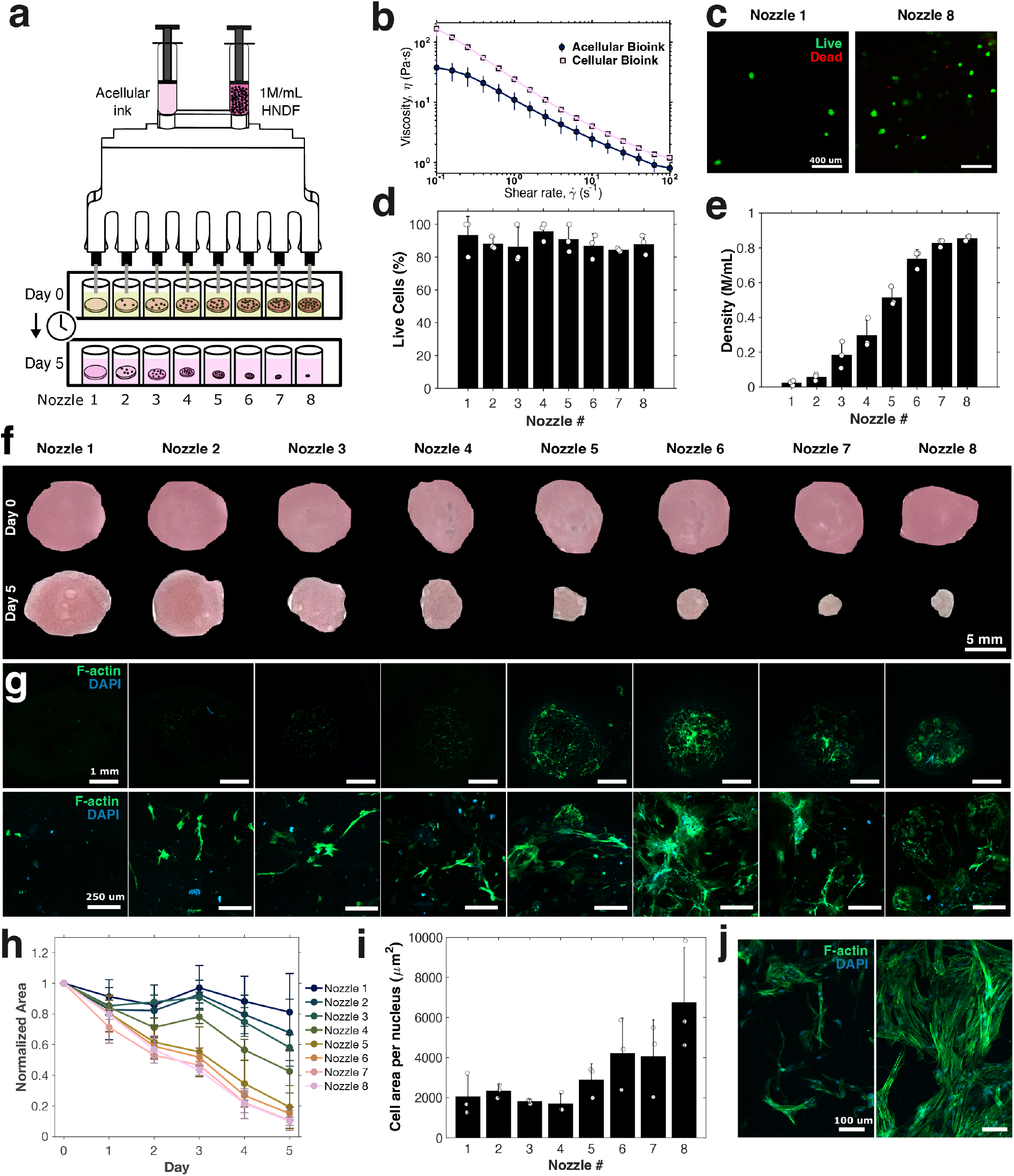
Bioprinting of tissue constructs with graded cell densities using a two-way gradient embedded multinozzle printhead. **a**, Schematic of the two-way gradient printhead illustrating printing of mixtures of acellular and HNDF-based bioinks and subsequent culturing for five days. **b**, Rheological characterization of acellular and HNDF-containing inks. **c–d**, Live–dead assay of bioprinted tissues showing representative images from nozzle 1 and nozzle 8 (**c**) and quantification of live-to-dead cell ratios across all eight nozzles (*n* = 3) (**d**). Scale bar, 400 *µ*m. **e**, Estimated cell density (million cells/mL) across all eight nozzles measured 1 h post-print (*n* = 3). **f**, Representative brightfield images of bioprinted tissues across all eight nozzles at day 0 (top) and day 5 (bottom). Scale bar, 5 mm. **g**, Confocal images at 2.5× (top) and 10× (bottom) magnification showing F-actin and DAPI staining of compacted tissues at day 5. Scale bars, 1 mm and 250 *µ*m. **h**, Change in tissue area from day 0 to day 5 across all nozzles (*n* = 3), with areas normalized to day 0 values. **i–j**, Changes in HNDF morphology quantified by cell area per nucleus (**i**) and visualized by representative high-resolution images showing thin, small, uncompacted HNDFs (left) and spindle-shaped, elongated, compacted HNDFs (right) (**j**). Scale bar, 100 *µ*m.

The resulting fibrin scaffolds exhibited high cell viability (approximately 90%) (**Fig. 3c–d**) and a gradient of fibroblast densities (**Fig. 3e, Supplementary Fig. 8b–d**). Consistent with prior findings^36^, scaffolds remained largely uncompacted below a critical density of approximately 0.5 million cells/mL (**Fig. 3f–h**). Below this threshold, HNDFs remained small and rounded, whereas above it they became spindle-shaped, elongated, and contractile (**Fig. 3i–j**). Notably, this experiment was performed in a single parallelized step, avoiding sequential formulation and printing steps that may be susceptible to time-dependent effects such as cell settling and ischemia.

### 2.4 GEM 3D Printing of Trileaflet Heart Valves

We next demonstrate the use of a three-way GEM printhead to optimize the composition of a photocurable hydrogel for 3D printed heart valves (**Fig. 4a**). Photocurable inks for embedded 3D printing must satisfy a set of competing constraints, including (i) appropriate rheology for printability and shape fidelity, (ii) minimal post-curing swelling, (iii) sufficient flexibility for valve opening, (iv) adequate stiffness and strength to prevent prolapse under back-pressure, and (v) high toughness to enable repeated cyclic loading. Satisfying these constraints requires careful tuning of multiple formulation parameters, including oligomer concentration, molecular weight, and multifunctional crosslinker content.

**Fig. 4:**
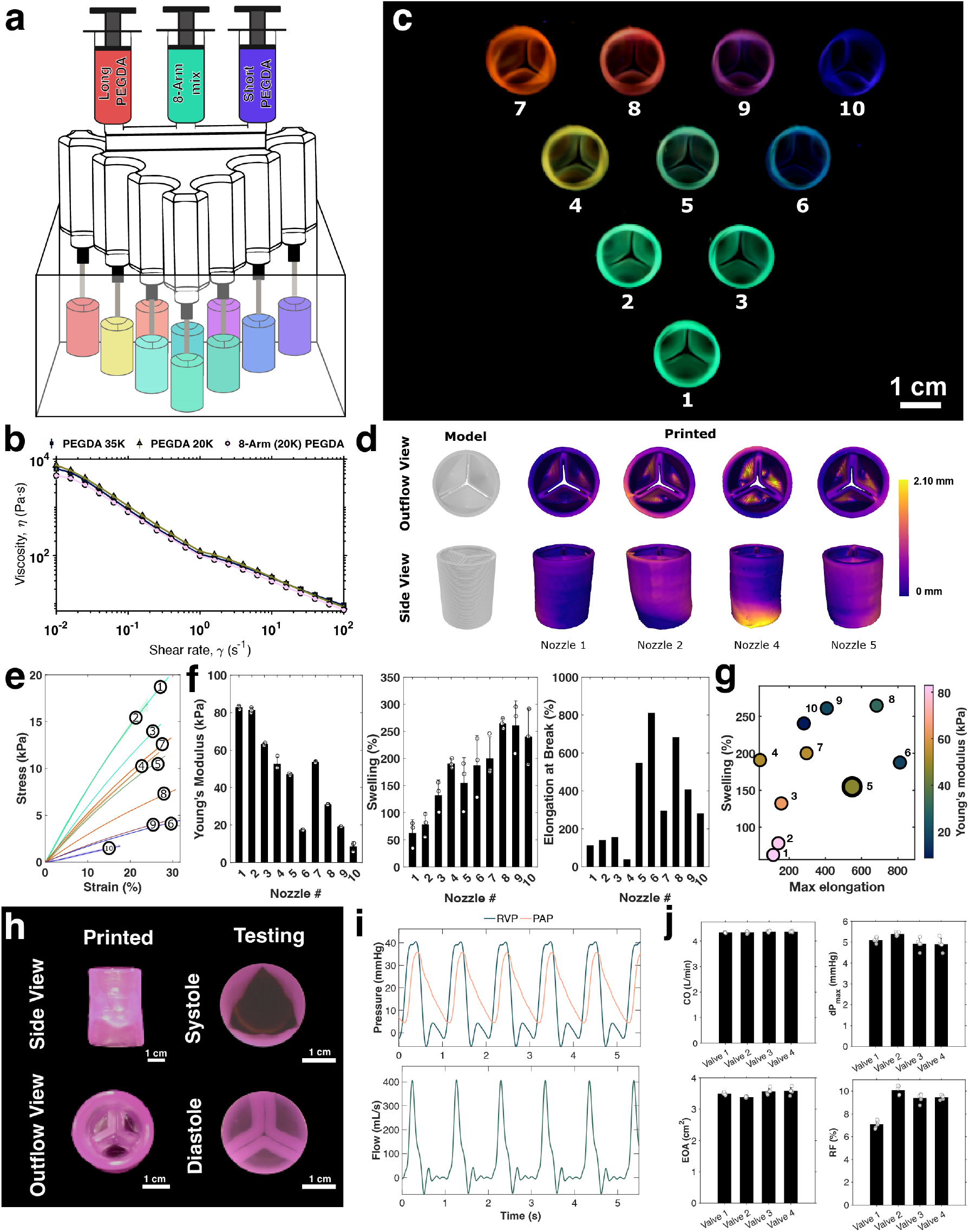
Trileaflet heart valve printing using a GEM printhead and material characterization. **a**, Schematic of a three-way gradient printhead illustrating parallel printing of an array of ten trileaflet heart valves. **b**, Viscosity as a function of shear rate for the three PEGDA-based input inks. **c**, Image of a printed array of ten valves showing graded polymer compositions, visualized using fluorescent microspheres. Scale bar, 1 cm. **d**, Micro-computed tomography (*µ*CT) analysis of valves printed from representative nozzle outputs (nozzles 1, 2, 4, and 5), shown in outflow and side views, with corresponding Hausdorff distance maps computed relative to the original stereolithography design. **e–g**, Material characterization of the ten nozzle outputs, including uniaxial tensile stress–strain curves (*n* = 3) (**e**), Young’s modulus (*n* = 3), swelling ratio (*n* = 3), and elongation to failure (*n* = 1) shown left to right (**f**), and a corresponding heatmap summarizing material properties across nozzle outputs, highlighting nozzle 5 as optimal for valve design and testing (**g**). **h**, Photography of a trileaflet valve post-printing (left; inflow and outflow views) and during hemodynamic testing at systole and diastole (right). Scale bar, 1 cm. **i–j**, Hemodynamic characterization of printed valves showing representative right ventricular pressure (RVP), pulmonary arterial pressure (PAP), and flow waveforms recorded over six consecutive cardiac cycles (**i**), and summary metrics calculated over eight to ten consecutive cycles for each tested valve (*n* = 4), including cardiac output (CO), maximum transvalvular pressure gradient (Δ*P*_max_), effective orifice area (EOA), and regurgitation fraction (RF) (**j**).

For example, increasing oligomer chain length can improve toughness^38^, but often increases swelling and reduces printability, whereas incorporation of multi-arm crosslinkers can suppress swelling at the expense of toughness^39^. These competing trade-offs motivate the use of GEM printheads to efficiently explore multi-dimensional formulation spaces.

We evaluated blends spanning low (4 kDa) to high (35 kDa) molecular weight linear poly(ethylene glycol) diacrylate (PEGDA), combined with varying fractions of an 8-arm PEGDA crosslinker. Two formulation families were investigated. The first family comprised (i) 10 wt% PEGDA 4 kDa, (ii) 10 wt% PEGDA 20 kDa, and (iii) a mixed formulation containing 2.5 wt% PEGDA 4 kDa, 2.5 wt% PEGDA 20 kDa, and 5 wt% PEGDA 8-arm (20 kDa). The second family comprised (i) 20 wt% PEGDA 20 kDa, (ii) 20 wt% PEGDA 35 kDa, and (iii) a mixed formulation containing 7.5 wt% PEGDA 20 kDa, 7.5 wt% PEGDA 35 kDa, and 5 wt% PEGDA 8-arm (20 kDa).

To improve printability and toughness, Carbopol was added as a rheological modifier at 0.3 wt/v% (ETD 2020) for the 4 kDa/20 kDa/8-arm formulations and at 0.75 wt/v% (Ultrez 20) for the 20 kDa/35 kDa/8-arm formulations, together with 0.1 wt% lithium phenyl-2,4,6-trimethylbenzoylphosphinate (LAP) photoinitiator. Within each formulation family, the three inks exhibited similar shear-thinning behavior (**Fig. 4b, Supplementary Fig. 9**), enabling consistent extrusion across outlets.

We first quantified the average polymer composition across the ten outlets of the three-way GEM printhead. Fluorescent microspheres (green, blue, and orange; 1–5 *µ*m diameter) were added to each inlet ink to validate the relative polymer fractions extruded from each outlet (**Supplementary Fig. 10**). Ten representative trileaflet valves printed in parallel are shown in **Fig. 4c**, demonstrating reproducible valve geometries across the outlet array.

To assess three-dimensional geometric fidelity, a triangular array of ten trileaflet valves was fabricated, and barium sulfate was incorporated into the 4 kDa/20 kDa/8-arm (20 kDa) PEGDA family to enable micro-computed tomography (*µ*CT) imaging (**Fig. 4d**). Quantitative analysis of a subset of valves revealed a mean root-mean-square (RMS) surface deviation of approximately 0.51 mm and a maximum Hausdorff distance of 2.10 mm, with deviations primarily localized to the valve root base and leaflet belly regions.

Achieving a suitably tough and flexible hydrogel is critical for valve function. To enable standardized mechanical testing independent of print pathing, inks were printed into dogbone molds and photocured. Tensile testing demonstrated that the 20 kDa/35 kDa/8-arm formulations produced hydrogels with superior mechanical performance compared to the 4 kDa/20 kDa/8-arm formulations (**Fig. 4e–f, Supplementary Fig. 11**). Across formulations, stiffness correlated primarily with the fraction of 8-arm PEGDA, whereas maximum extensibility and deformation were dominated by the higher molecular weight PEGDA components.

Post-photocuring swelling was quantified by measuring dimensional changes in printed hydrogel samples. Increased concentrations of PEGDA 8-arm reduced swelling, whereas higher fractions of PEGDA 20 kDa and 35 kDa increased swelling (**Fig. 4f**). Among the tested compositions, the formulation corresponding to outlet nozzle 5 (5.4% PEGDA 20 kDa, 12.1% PEGDA 35 kDa, and 2.5% PEGDA 8-arm (20 kDa)) achieved an optimal balance of mechanical strength and swelling behavior (**Fig. 4g**). This formulation was independently prepared for single-nozzle printing of valves for hemodynamic testing.

Following printing and photocuring (**Fig. 4h**), four valves were individually integrated into a univentricular flow simulator for hemodynamic evaluation^40^ (**Supplementary Fig. 12**). Under physiological pulmonary valve loading conditions, the printed valves exhibited appropriate opening and closure, as shown by co-registered images at peak systole and end diastole (**Fig. 4i; Supplementary Video 2**). Representative right ventricular and pulmonary artery pressure traces, together with pulmonary artery flow waveforms recorded over five consecutive cardiac cycles, demonstrated proper leaflet motion and unobstructed antegrade flow.

At a cardiac output of approximately 4.5 L/min (heart rate ∼ 60 bpm, stroke volume ∼ 75 mL), valves exhibited a maximum transvalvular pressure gradient of 5.1 ± 0.2 mmHg, an effective orifice area of 3.5 ± 0.1 cm^2^, and a regurgitant fraction of 9.0 ± 1.3% (**Fig. 4j**), indicating appropriate systolic and diastolic performance. Compared with previously reported PEGDA-based valves^19^, this optimized formulation achieved a 106% increase in effective orifice area and reductions of 52% and 68% in maximum pressure gradient and regurgitation, respectively. All pressure and flow waveforms, together with heart rate and stroke volume inputs, are provided in **Supplementary Fig. 13**.

## 3 Conclusions

We have demonstrated two-way, three-way, and four-way gradient embedded multinozzle (GEM) print-heads capable of fabricating arrays of three-dimensional constructs with systematically varied material compositions. By tuning mixer design, GEM printheads can generate linear or sigmoidal composition gradients, enable high-viability cell printing, and resolve critical cell-density thresholds for scaffold compaction. We further applied a three-way GEM printhead to optimize PEGDA-based formulations for printing tough, trileaflet hydrogel heart valves. More broadly, the ability of GEM printheads to rapidly generate compositionally graded, cell-laden constructs in arrayed formats may enable scalable platforms for in vitro drug screening and dose–response studies in engineered tissues. Beyond bioprinting, GEM printheads may find broader applications in the manufacturing of graded metamaterial structures^41^ or graded robotic actuators^42^.

GEM printheads have several limitations. The distributed channel networks required for mixing and redistribution introduce increased dead volume and hydraulic resistance, particularly at larger length scales. For low-viscosity but costly inks (e.g., bioinks), channel dimensions may be reduced within the resolution limits of stereolithographic printing to minimize material loss. Conversely, for low-cost, high-viscosity inks, wider channels can reduce hydraulic resistance at the expense of increased dead volume. For highly viscous materials, passive mixing alone may be insufficient, necessitating the incorporation of active mixing strategies. While the present GEM printhead designs assume similar ink rheologies, future asymmetric channel architectures may enable mixing of inks with differing rheological properties.

GEM printheads are distributed as variable-driven design files to enable rapid adaptation of nozzle architecture to application-specific scale and throughput requirements. By parallelizing formulation, mixing, and printing, GEM printheads provide a general strategy to accelerate exploration and optimization of inks for direct ink writing of functional constructs.

## 4 Methods

### Nozzle Design and Manufacturing

All nozzles were designed in OnShape (v. 1.208.68980.736ce7a5e6f5, PTC Inc.). Nozzles were printed on Formlabs Form 4 or Form 3B printers (Formlabs) using Clear V5 (FLGCPCL05), Clear V4 (FLGCPCL04), or Clear V4.1 (FLGCPCL041) resin with a 0.100 mm layer height. For bioprinting, nozzles were printed with Biomed Black Resin (Formlabs, FLBMBL01) on a Formlabs Form 3B with a 0.050 mm layer height.

Nozzles were submerged in a FormWash IPA rinse for 30 min after printing, then channels were flushed with 30 mL BD Hamilton Luer-Lock syringes (BD, SKU: 302832; GTIN: 00382903028320) using fresh isopropyl alcohol (MaxTite, isopropyl alcohol 99.9%). Channels were blow-dried with air to remove alcohol and resin residue and then allowed to dry for 3 days at room temperature on paper towels (Scotts Shop Towel, U-Line, S-15737) covered in aluminum foil (Reynolds) to protect from light. After drying, nozzles were cured in a Formlabs FormCure at 60 °C for 15 min (Clear V4, V4.1, and V5 resins) or 70 °C for 60 min (Biomed Black). After curing, nozzles were kept dry at room temperature until use. To ensure no channels were clogged, each nozzle was flushed with Milli-Q water using 30 mL BD Hamilton Luer-Lock syringes prior to use.

### Nozzle Photography

To improve transparency, nozzles were coated in Sylgard 184 base (Dow 4019862) and photographed using a Canon EOS 5D Mark IV camera with a Canon Macro Lens EF 180 mm f/3.5 Ultrasonic Stabilizer, Canon EF 24–105 mm lens, or Laowa 24 mm f/14 2× lens (Venus Optics) for videography. UV lamps (405 nm; Geetech, Amazon B0DCNV29LP) were used to illuminate fluorescent pigments. Mirrors (Bright-Creations, Amazon B07T8Y58KR) were used as needed to adjust viewing angles for videography.

### Base Ink Preparation

Carbopol (ETD 2020, Lubrizol) was dissolved in Milli-Q water at 0.3 wt% overnight using an overhead impeller (Fisherbrand CompactDigital Overhead Stirrer, Fisher Scientific, 14500211) at 685–950 rpm with a viscoelastic stirrer (McMaster-Carr, 35325K57), then titrated to pH 7 using 10 M NaOH (Thermo A16037.36). After titration, the solution was stirred overnight at room temperature at 100 rpm to remove bubbles.

Either 0.02 wt% fluorescent microspheres (Cospheres FMO, FMOR, FMB, FMG, or FMV) or 1 wt% thermoplastic neon fluorescent pigment (TechnoGlow Magenta UVP-MAG-E001Z, Blue UVP-BLU-E001Z, Orange UVP-FRO-E001Z, or Yellow UVP-YLW-E001Z) was added to 30 g of ink in FlackTek cups (FlackTek, SC 90) and mixed in a FlackTek SpeedMixer (330-100 PRO) at 2000 rpm for 2 min to homogenize. Inks were then loaded into 30 mL BD Hamilton Luer-Lock syringes (BD SKU: 302832; GTIN: 00382903028320) fitted with end tips (Qosina, 61726) and centrifuged at 300*×g* for 5 min (Thermo ST 16 Sorvall). Plungers were loaded from the back alongside a stainless steel wire (Youngtiger 304 Stainless, X002BBCQJ5) to allow trapped air to escape, after which the wire was removed. Inks were stored at room temperature for up to 2 days before use.

### 20k/35k/8-arm Valve Parent Ink Preparation

Carbopol Ultrez 20 (Lubrizol) was prepared at 3 wt% using the protocol above. PEGDA 20k (Poly-sciences, SKU: 26280), PEGDA 35k (Polysciences, SKU: 263423), and 8-arm (20k) (tripentaerythritol) PEGDA (JenKem Technology, SKU: 8ARM(TP)-ACLT) were dissolved at 40 wt% in Milli-Q water overnight on a roller mixer (Roller Mixer TRM, 8011560) at 4 °C in 50 mL Falcon tubes (Sigma, CLS352070). LAP (Sigma 900889) was dissolved at 5 wt/v% in Milli-Q water overnight on a roller mixer at 4 °C.

PEGDA 20k and PEGDA 35k parent inks were prepared by combining 4.60 mL Milli-Q water, 5 g of 3 wt% Carbopol Ultrez 20, 10 mL of 40 wt/v% PEGDA solution (20k or 35k), 400 *µ*L of 5 wt/v% LAP, and 0.02 wt% fluorescent microspheres (Cospheres FMB for PEGDA 20k; FMO for PEGDA 35k). The 8-arm (20k) PEGDA parent ink was prepared by combining 4.60 mL Milli-Q water, 5 g of 3 wt% Carbopol Ultrez 20, 2.5 mL of 40 wt/v% 8-arm (20k) PEGDA, 3.75 mL of 40 wt/v% PEGDA 20k, 3.75 mL of 40 wt/v% PEGDA 35k, 400 *µ*L of 5 wt/v% LAP, and 0.02 wt% microspheres (Cospheres FMG).

Each parent ink was mixed in an SC 90 cup using the SpeedMixer at 2000 rpm for 2 min, loaded into 30 mL BD Hamilton Luer-Lock syringes with end tips, and centrifuged at 300*×g* for 5 min. Plungers were inserted using the stainless-steel wire method described above. Inks were stored at 4 °C protected from light (aluminum foil) and used within 1 day.

### 4k/20k/8-arm Valve Parent Ink Preparation

Carbopol ETD 2020 (Lubrizol) was prepared at 0.6 wt% using the protocol above. PEGDA 20k (Polysciences, SKU: 26280), PEGDA 4k (Polysciences, SKU: 15246), and 8-arm (20k) (tripentaerythritol) PEGDA (JenKem Technology, SKU: 8ARM(TP)-ACLT) were dissolved at 20 wt% (PEGDA 4k and PEGDA 20k) and 40 wt% (8-arm (20k)) in Milli-Q water overnight at 4 °C on a roller mixer in 50 mL Falcon tubes. LAP was prepared at 5 wt/v% similarly.

PEGDA 4k and PEGDA 20k parent inks were prepared by combining 15 g of 0.6 wt% Carbopol ETD 2020, 15 mL of 20 wt/v% PEGDA solution (4k or 20k), 600 *µ*L of 5 wt/v% LAP, and 0.02 wt% fluorescent microspheres (Cospheres FMB for PEGDA 20k; FMO for PEGDA 4k). The 8-arm (20k) PEGDA parent ink was prepared by combining 18.75 g of 0.6 wt% Carbopol ETD 2020, 3.75 mL of 40 wt/v% 8-arm (20k) PEGDA, 3.75 mL of 20 wt/v% PEGDA 20k, 3.75 mL of 20 wt/v% PEGDA 35k, 600 *µ*L of 5 wt/v% LAP, and 0.02 wt% microspheres (Cospheres FMG). Each ink was SpeedMixer-homogenized (2000 rpm, 2 min), syringe-loaded, and centrifuged at 300*×g* for 5 min, then stored at 4 °C protected from light and used within 1 day.

For *µ*CT characterization, BaSO_4_ (Thermo, 222510050) was added to each parent ink at 20 wt/v% prior to mixing; SpeedMixer, centrifugation, and loading steps were performed identically.

### Embedded Printing Bath Preparation

Carbopol ETD 2020 was prepared at 0.3 wt% as described above, then diluted with an equal volume of Milli-Q water to reach 0.15 wt%. The diluted Carbopol was stirred overnight with an overhead impeller to remove bubbles and stored at 4 °C covered with Saran wrap to prevent evaporation. If bubbles remained, the bath was degassed by centrifugation in 500 mL conicals (Thermo, 332270) at 2000*×g* for 5 min.

### Extrusion Printing

Each nozzle was fitted with Nordson EFD tips (ID 0.41 mm; length 0.25–1.5 in; Nordson 7018260, 7018272, 7018263, 7018266). The same tip length was used across nozzle outputs. Parent inks were mounted onto a custom-built 3D printer derived from the low-cost bioprinter design described by Weiss et al.^19^. An Octopus MAX EZ V1.0 BTT controller (Bigtree Tech) with EZ2209 TMC drivers was used to create a 10-axis system (four extrusion axes, four Z-axes, and X/Y motion). The frame was machined from aluminum (McMaster-Carr 9057K36 and 9057K38) on a Haas VF 2-SS CNC with toolpaths generated in Fusion 360 CAM (v. 2605.1.52). Each extrusion axis used a 5:1 planetary gear stepper motor (StepperOnline CN-11HS20-0714S-PG15) coupled to a screw axis (McMaster-Carr 93675A498) via a shaft collar (McMaster-Carr 5395T111). The screw axis was flat-ground (KEF PSD 10, 1022024) to improve coupling strength, and a shaft collar housing was 3D printed (Markforged Black Onyx) on a Markforged MarkTwo printer (assembly instructions in Supplementary Fig. 2). Firmware was modified so that all four extrusion axes moved together, and all four Z-axes moved together, yielding effective single-head X/Y/Z motion. The printer was mounted on vibration absorbers (McMaster-Carr 9232K14).

For photocurable prints, parent inks, nozzle, and printer were protected from UV exposure. After printing, baths were illuminated with 405 nm light for 15 min to crosslink.

### Pressure-Based Printing

Pressure-based extrusion was performed on a custom Aerotech gantry system (Automation 1 software; XY ABCD axes driven by XR3 cards; additional axes via XC2 cards). A Nordson EFD Ultimus V dispenser (Nordson 7012590) was used with a custom pressure connector formed from three single-syringe connectors (McMaster-Carr 66275A14) and two wye connectors (McMaster-Carr 53055K507). Custom clamps and syringe holders were 3D printed (Markforged Black Onyx) to mount syringes (McMaster-Carr 66305A54). Valve-printing code from Weiss et al.^19^ was adapted, and custom acrylic boxes were fabricated from acrylic sheets (McMaster-Carr 8560K239) by laser cutting (Glowforge 60 W) and solvent bonding (Weld-On, US Plastic 46871).

### Spectral Imaging

After inks were passed through each nozzle and collected at the tips in 1 mL microcentrifuge tubes, samples were centrifuged at 2000*×g* for 2 min. Aliquots of 100 *µ*L (n=3) from each nozzle output and 100 *µ*L from parent inks were dispensed into a 96-well plate (Corning 3368). Plates were scanned from the bottom using a Tecan Spark multimode plate reader. Wells were excited at 2–4 wavelengths depending on the number of parents (bandwidth 10 nm): 405 nm (FMB), 488 nm (FMG), 561 nm (FMO), and 584 nm (FMV). Emission spectra were recorded at 425–470 nm (FMB), 515–540 nm (FMG), 610–758 nm (FMO), and 650–800 nm (FMV) in 5 nm steps, using gain 100 for FMO/FMB/FMV and gain 75 for FMG, with 30 flashes per well. Parent ink spectra were used to generate fingerprint spectra, and spectral unmixing was performed by stacking spectra into a column matrix and solving for component mixtures via linear inversion.

### Confocal Imaging

The 96-well plate was imaged on a ZEISS LSM 980 confocal microscope using Zen software (Zen 3.8, ZEISS) with settings matched to the plate reader. Wells were excited at 405, 488, and 561 nm, with emission bands set to 425–470 nm, 515–540 nm, and 610–800 nm to minimize cross-talk. Z-stacks were acquired in full-stack, per-track mode at 1024×1024 resolution, frame time ∼1.0 s, 2× averaging, and 16-bit depth, at 2.5× and 10× magnifications (n=3). Confocal imaging was not performed for four-way nozzles due to cross-talk between FMO and FMV beads. Images were analyzed in FIJI (Java 21.0.7, 64-bit) and custom MATLAB scripts that split channels, computed maximum-intensity projections, and segmented particles to quantify bead counts. Control samples were used to establish baselines for unbiased fractional composition.

### Rheological Sample Preparation

Rheology samples were prepared similarly, with total volume limited to 500 *µ*L. Samples were prepared in 1 mL microcentrifuge tubes (Fisher 02-682-002) using positive-displacement pipettes (Gilson Micro-man FD10004/FD10005/FD10006) and capillary pistons (Fisher F148560G/F148414G/F148714G). For PEGDA 20k and PEGDA 35k parent inks, samples were prepared by combining 120 *µ*L Milli-Q water, 125 *µ*L of 3 wt% Carbopol Ultrez 20, 250 *µ*L of 40 wt/v% polymer solution, and 10 *µ*L of 5 wt/v% LAP, then pipetting up and down 50 times. The 8-arm (20k) PEGDA sample was prepared by combining 120 *µ*L Milli-Q water, 125 *µ*L of 3 wt% Carbopol Ultrez 20, 63 *µ*L of 40 wt/v% 8-arm (20k) PEGDA, 94 *µ*L of 40 wt/v% PEGDA 20k, 94 *µ*L of 40 wt/v% PEGDA 35k, and 10 *µ*L of 5 wt/v% LAP, then pipetting 25 times. For ETD 2020 (0.3 wt%), a 500 *µ*L sample was collected during ink preparation. For fibrin ink rheology, 1 mL samples of acellular and cellular bioinks were prepared as described below.

### Rheological Characterization

Rheology of hydrogel inks (0.3 wt% ETD and 4k/20k/8-arm (20k) PEGDA inks) was measured on a stress-controlled rheometer (TA Instruments AR-G2) using a 25 mm parallel-plate geometry and Peltier temperature control at 25 ± 0.01 °C with a 0.5 mm gap. For 20k/35k/8-arm (20k) PEGDA inks, an 8 mm serrated parallel-plate geometry was used with a 0.5 mm gap at 25 ± 0.01 °C. For fibrin inks, a TA Discovery Hybrid Rheometer HR-2 was used with crosshatched 20 mm plates at 25 ± 0.01 °C and a 0.60 mm gap. Samples were loaded, trimmed, and subjected to oscillatory shear over 100–0.1 rad s^*−*1^. Storage (*G*^*′*^) and loss (*G*^*′′*^) moduli were measured with frequency set to 0.5 Hz while varying oscillatory strain amplitude from 1000% to 0.1%.

### Flow Characterization

Flow uniformity across nozzle outlets was quantified by measuring the mass extruded from each outlet for two-, three-, and four-way printheads using 0.3 wt% ETD 2020 ink. Approximately 1 g was extruded per nozzle per trial. Extrusions were collected into 2 mL low-retention snap-cap microcentrifuge tubes (Thermo Fisher Scientific, Cat. No. 3434). Each condition was repeated three times (n=3). Mass was measured using an analytical balance (Mettler Toledo, XSR 104).

### Micro-CT Scanning

Valves were printed into cylindrical custom containers (Formlabs Form 3B, Clear V4 resin) and immediately scanned using a Bruker SkyScan 1276 CT scanner (85 kV, 200 *µ*A; rotation step 0.400 degrees; Al 1 mm filter; 2-frame averaging). Reconstruction was performed using NRecon (Micro Photonics). STLs were generated in 3D Slicer (v. 5.6.2) and Hausdorff distance was computed in MeshLab (v. 2023.12).

### Cell Culture

Cells were maintained at 37 °C with 5% CO_2_ in T75 or T175 flasks (Thermo, 156800 and 159926). Human neonatal dermal fibroblasts (HNDFs; Lonza, CC-2509) were cultured in DMEM (Thermo, 11965092) supplemented with 10% FBS (Sigma, F4135) and 1% Penicillin-Streptomycin (5000 U/mL, Fisher, 15070063). Media was changed every 3–5 days and cells were passaged 1:5 upon confluence.

### Bioink Preparation

Gelatin solution was prepared by dissolving type B gelatin (Fisher, G7-500) at 15% w/v in PBS containing calcium and magnesium (Thermo Gibco, 14040141), stirring at 60 °C for 6 h, adjusting to pH ∼7.3 with 1 M NaOH, and vacuum sterile filtration. Bovine fibrinogen (EMD Millipore, 341573) was dissolved at 50 mg/mL in DMEM at 37 °C for ∼2 h. Gelbrin bioink was prepared by mixing fibrinogen (33 mg/mL final) and gelatin (5% w/v final) from 50 mg/mL fibrinogen and 15% gelatin stocks.

HNDFs were passaged using TrypLE (Thermo Gibco, 12604013) for 3 min, centrifuged at 300×*g* for 5 min (Thermo Sorvall X-Pro Series), and resuspended directly into fibrinogen before mixing with gelatin to form 5 mL final cellular bioink. An acellular ink of identical composition was prepared in parallel. FRESH 2.0 support bath was prepared per published protocol^37^, washed in DMEM, and supplemented with 10 U/mL thrombin (Sigma-Aldrich, T4648). The GEM printhead was sterilized by flushing with 70% ethanol (from GoldShield 200-proof ethanol) and 0.1 M NaOH, followed by extensive PBS rinsing. Inlets were pre-filled with PBS and bubbles were avoided during flushing.

### 3D Printing of Bioinks

Cellular and acellular bioinks were loaded into separate 5 mL luer-lok syringes (BD, SKU: 309646; GTIN: 00382903096466) fitted with 1-inch, 0.33 mm ID tips (Nordson EFD Orange 7018305). Syringes were attached to the printhead while pressing tips into a sterile PDMS slab (Sylgard 184, 10:1) to prevent leakage and air ingress. Constructs were printed into the FRESH bath, stabilized at room temperature for 10 min, and then incubated at 37 °C to melt the support.

### Tissue Culture and Endpoint Processing

Constructs were maintained in DMEM-based medium containing 1% Pen/Strep, 10% FBS, and 0.4 TIU/mL aprotinin (GoldBio, A-655-25) with full-volume daily media changes in six-well plates. Samples were fixed in 4% paraformaldehyde (Thermo, AA47392-9L). Cultures were terminated 1 h post-print for live–dead analysis (n=3) and cell density measurements (n=3), and on day 5 for compaction assays (n=3). Calcein-AM and ethidium homodimer-1 (Thermo Fisher Scientific, L3224) were used for live–dead assays. Phalloidin (488 nm; Thermo Scientific A12379) and nuclei markers (Hoechst, ApexBio A3472) were used for other assays.

### Confocal Imaging of Bioprinted Samples

Samples were imaged on a ZEISS LSM 980 confocal microscope using Zen software (Zen 3.8, ZEISS). Excitation was performed using 405, 488, and 561 nm lasers. Z-stacks were acquired in full-stack, per-track mode at 1024×1024 resolution, frame time *∼*2.0 s, and 4× averaging. Images were acquired at 2.5× and 10× magnifications.

### Tensile Testing

Uniaxial tensile testing was performed using a dynamic mechanical analyzer (ARES-G2, TA Instruments). Each nozzle outlet was collected, centrifuged, and dispensed into custom acrylic molds using positive-displacement pipettes, then UV-cured under 405 nm light for 5 min. Specimens were removed using a flat-backed spatula (Fisher, S41699) and stored overnight at 4 °C on water-spritzed paper towels wrapped in Saran wrap to prevent dehydration. Specimens were tested with 25 mm gauge length, 6.5 mm width, and 3.18 mm thickness (1/8 in). Samples were pulled at 0.25 mm/s with acquisition at 10 Hz. Young’s modulus was calculated from the 0.5–5% linear region of the stress–strain curve.

### Swelling Analysis

Disks were collected using a disposable 4 mm biopsy punch from material extruded into 1/8-inch-deep molds. For each of the ten outputs of the three-way nozzle, *n* = 3 disks were incubated in 1× PBS (Thermo, 18912014) overnight. Images were acquired at baseline and after 24 h. Disk area was measured in ImageJ (Java 1.8.0 345, 64-bit) and swelling ratio was computed relative to baseline.

### Valve Design

The valvular leaflet geometry was generated according according to Hamid et. al^43^ and reported previously in Weiss et. al^19^. First, an elliptic paraboloid centered at the valvular root’s edge was generated, and paraboloid points on the outside of the root geometry were omitted. A subsegment of the valvular root was then created by two planes intersecting the centerline of the valvular root and separated by 60 degrees in each direction from the center of the paraboloid. All the leaflet points that crossed the generated planes were modified to be on the respective plane with an offset factor added to prevent the leaflets from overlapping and curing together during printing. Finally, the generated leaflet points were symmetrically patterned in a circular manner around the centerline of the valvular root.

### Valve Printing

An array of ten valves was printed using the three-way printhead with inks composed of 10% PEGDA 4k, 10% PEGDA 20k, and 5% 8-arm PEGDA, and 7.5% PEGDA 20k and 7.5% PEGDA 35k as described above. Valves selected for functional testing were printed using a single nozzle after optimization through mechanical and swelling analyses. These valves used a formulation of 5.4% PEGDA 20k, 12.1% PEGDA 35k, and 2.5% 8-arm PEGDA.

A master mixture was prepared by combining 6.02 mL of PEGDA 20k (40 wt/v%) solution, 13.67 mL of PEGDA 35k (40 wt/v%) solution, 2.81 mL of 8-arm (20k) PEGDA (40 wt/v%) solution, 11.25 g of 3 wt% Ultrez 20 solution, 0.9 mL of LAP (5 wt/v%) solution, 10.35 mL of Milli-Q water, and 0.02 wt/v% fluorescent microspheres (Cospheres FMV). The mixture was SpeedMixer-homogenized (SC 90 cup, 2000 rpm, 2 min), loaded into 10 mL BD Luer-Lok syringes (SKU: 302995; GTIN: 00382903029952) with Luer-Lok end tips (Qosina, 61726), and centrifuged at 2000 ×*g* for 2 min (Thermo Sorvall ST 16).

Plungers were inserted using the steel-wire method, and syringes were stored at 4 °C protected from light. Single-nozzle valve-printing code was adapted from Weiss et al.^19^. Following printing, valves were UV-cured under 405 nm illumination for 10 min, rinsed with DI water, and stored dry for up to 48 h before testing.

### Valve Testing and Hemodynamic Analysis

Valves were evaluated in a univentricular ex vivo heart simulator (SuperPump, ViVitro Labs)^40^. Valves were mounted using a 34 mm woven vascular graft as reinforcement. At the basal region, the valve was secured with three mattress sutures using 4-0 polypropylene. To minimize the risk of material tearing, the graft was wrapped circumferentially around the valve below the leaflet nadir. To ensure a fluid-tight seal, the graft–valve interface was reinforced with 0 silk sutures. At the apical region, the graft was shaped to conform to the top-mount geometry using purse-string sutures with 2-0 polyester. The graft was then wrapped around the valve above the commissures and secured with 0 silk sutures to prevent leakage.

The simulator recreates pulmonary/systemic pressure and flow profiles using saline, with peripheral resistance, stroke volume, and heart rate controlled using ViVitest software. Settings were tuned to approximate physiological right-heart conditions (target cardiac output *≈* 4.5 L/min; mean right ventricular pressure *≈* 20–30 mmHg).

Pulmonary flow was measured using 25 mm ultrasonic flow probes (Carolina Medical Electronics) and pressures were measured using ventricular and pulmonary artery transducers (Utah Medical Products). Sensors were zeroed to ambient pressure prior to each trial and flow probes were recalibrated to ensure a no-flow baseline. High-speed valve motion was recorded from the outflow side at 1057 fps and 1280×1024 resolution (Chronos 1.4; Kron Technologies). Hemodynamic metrics were computed in MATLAB R2024b using established analysis scripts^44^. Right ventricular pressure, pulmonary artery pressure, and pulmonary artery flow were imported directly from the simulator output. Stroke volume, cardiac output, transvalvular pressure gradient, effective orifice area, and regurgitant fraction were calculated over 8–10 consecutive cardiac cycles.

## Acknowledgements

L.R., S.S., and J.D.W. contributed equally to the manuscript. L.R. was funded by a Burroughs Wellcome Fund Career Awards at the Scientific Interface, S.S. was funded by the National Science Foundation Graduate Research Fellowship Program under award number DGE-1656518 and by the Stanford Graduate Fellowship. J.D.W. was funded by the National Science Foundation Graduate Research Fellowship Program under award number DGE-1656518 and Stanford Bio-X. C.M.W was funded by the National Science Foundation Graduate Research Fellowship Program under award number DGE-2146755. Research reported in this publication was funded by the National Heart, Lung, and Blood Institute of the National Institutes of Health under award number DP2HL168563. The content is solely the responsibility of the authors and does not necessarily represent the official views of the National Institutes of Health.

## Author Contributions

Conceptualization: L.R., S.S., J.D.W., and M.A.S.-S. Methodology: L.R., S.S., J.D.W., S.H., F.S.S., A.S., M.S., J.D., K.M., D.R.R., Z.C.R., Q.W., C.M.W., J.H.K., E.A.A., M.M., and M.A.S.-S. Formal analysis: L.R., S.S., J.D.W., and M.A.S.-S. Visualization: L.R., S.S., J.D.W., and F.S.S. Validation: L.R., S.S., J.D.W., and S.H. Investigation: L.R., S.S., J.D.W.,S.H., F.S.S., M.S., A.S, A.R. and M.A.S.-S.. Software: T.R., T.T., Z.C.R., C.M.W., and J.H.K. Cell work: L.R., J.D.W, K.M.,D.R.R. and Q.W. Theoretical Derivations: S.S., J.D., J.D.W. Resources: E.A.A., M.M., and M.A.S.-S. Data curation: L.R., S.S., and M.A.S.-S. Funding Acquisition: M.M., E.A.A., and M.A.S.-S. Writing: L.R., S.S., J.D.W., and M.A.S.-S., and all authors contributed to reviewing and editing the manuscript.

