## Supplementary Information for "Gradient Multinozzle 3D Printing"

### List of Contents

**Supplementary Section 1** – GEM theory
**Supplementary Figure 1** – Construction of GEM nozzles using variable-driven CAD
**Supplementary Figure 2** – High-torque stepper motor adapter assembly
**Supplementary Figure 3** – Confocal imaging and processing; flow output for each GEM; rheological characterization
**Supplementary Figure 4** – Theory of GEM
**Supplementary Figure 5** – Confocal imaging characterization of two- and three-way printheads
**Supplementary Figure 6** – Spectral unmixing characterization of four-way printheads and two-way imperfect mixing
**Supplementary Figure 7** – Effect of ink rheology on two-way printhead performance
**Supplementary Figure 8** – Extended fibrin rheology and confocal imaging of cell printing
**Supplementary Figure 9** – Rheology of 10% PEGDA and 20% PEGDA formulations
**Supplementary Figure 10** – Confocal imaging of valve outputs
**Supplementary Figure 11** – Dynamic mechanical analysis of PEGDA blends and valve hemodynamics
**Supplementary Video 1** – 3D gradient multinozzle embedded printing of trileaflet valves
**Supplementary Video 2** – Hemodynamic testing of trileaflet valves

### 1 Mixer graph construction

We represent each gradient mixer architecture as a directed acyclic graph (DAG)

$$G = (V, E),$$

where nodes $V$ represent mixing sites and edges $E$ represent channels between sites. Nodes are organized into $\mathcal{L}$ discrete layers indexed by

$$\ell \in \{1, 2, \ldots, \mathcal{L}\},$$

with $\ell = 1$ containing the inlets and $\ell = \mathcal{L}$ containing the outlets. Let $V_\ell \subset V$ denote the set of nodes in layer $\ell$, and let $E_{\ell\to\ell+1} \subset E$ denote edges connecting layer $\ell$ to $\ell + 1$. The complete topology is determined by $\{V_\ell\}_{\ell=1}^{\mathcal{L}}$ and $\{E_{\ell\to\ell+1}\}_{\ell=1}^{\mathcal{L}-1}$.

Throughout, let $n_{\text{in}} \in \{2, 3, 4\}$ denote the number of inlets and let $k := |V_{\mathcal{L}}|$ denote the number of outlets. See **Supplementary Fig. 4** for subsequent diagram.

#### 1.1 Two-input (binary strip) architecture ($n_{\text{in}} = 2$)

The layer sizes satisfy

$$|V_\ell| = \ell + 1, \qquad \ell = 1, \ldots, \mathcal{L},$$

so $|V_1| = 2$ and $k = |V_{\mathcal{L}}| = \mathcal{L} + 1$. Label the nodes in layer $\ell$ by $\{u_1, \ldots, u_{|V_\ell|}\}$ and in layer $\ell + 1$ by $\{v_1, \ldots, v_{|V_{\ell+1}|}\}$. Each node connects to two neighbors in the next layer:

$$u_i \longrightarrow \{v_i, v_{i+1}\}, \qquad i = 1, \ldots, |V_\ell|.$$

Thus,

$$E_{\ell \to \ell+1} = \big\{(u_i \to v_i),\ (u_i \to v_{i+1}) \;\big|\; i = 1, \ldots, |V_\ell|\big\}.$$

#### 1.2 Three-input (nested triangle) architecture ($n_{\text{in}} = 3$)

Layer $\ell$ forms a triangular lattice with

$$|V_\ell| = \frac{(\ell+1)(\ell+2)}{2}, \qquad \ell = 1, \ldots, \mathcal{L},$$

so $|V_1| = 3$ and $k = |V_{\mathcal{L}}| = \frac{(\mathcal{L}+1)(\mathcal{L}+2)}{2}$. Index nodes by $(r, c)$ and write $\mathrm{triID}_\ell(r, c)$ for row $r$, column $c$ with

$$1 \le r \le \ell + 1, \qquad 1 \le c \le r.$$

Each node has up to three children:

$$(r, c) \longrightarrow \{(r, c), (r+1, c), (r+1, c+1)\}.$$

Equivalently, for all valid $(r, c)$,

$$\mathrm{triID}_\ell(r,c) \to \mathrm{triID}_{\ell+1}(r,c), \quad \mathrm{triID}_\ell(r,c) \to \mathrm{triID}_{\ell+1}(r+1,c), \quad \mathrm{triID}_\ell(r,c) \to \mathrm{triID}_{\ell+1}(r+1,c+1).$$

#### 1.3 Four-input (square pyramidal) architecture ($n_{\text{in}} = 4$)

Layer $\ell$ forms a square grid with

$$|V_\ell| = (\ell+1)^2, \qquad \ell = 1, \ldots, \mathcal{L},$$

so $|V_1| = 4$ and $k = |V_{\mathcal{L}}| = (\mathcal{L}+1)^2$. Let $\mathrm{gridID}_\ell(i, j)$ denote the node at $(i, j)$ with $1 \le i, j \le \ell + 1$. Each node connects to four children:

$$(i, j) \longrightarrow \{(i, j), (i+1, j), (i, j+1), (i+1, j+1)\}.$$

##### 1.4 Layer sizes and branching patterns

| $n_{\text{in}}$ | Layer shape | $\lvert V_\ell\rvert$ | Children per parent |
|---|---|---|---|
| 2 | 1D strip | $\ell+1$ | 2 |
| 3 | 2D triangle | $\frac{(\ell+1)(\ell+2)}{2}$ | 3 |
| 4 | 2D square | $(\ell+1)^2$ | 4 |

Edges only connect adjacent layers. In particular, if $(u \to v) \in E$, then $u \in V_\ell$ and $v \in V_{\ell+1}$ for some $\ell$, and the graph is acyclic.

#### 2 Geometric conventions and connector-volume constants

Nodes are placed at the center of a regular connector geometry: (i) the midpoint of a line segment for $n_{\text{in}} = 2$, (ii) the centroid of an equilateral triangle for $n_{\text{in}} = 3$, and (iii) the center of a square for $n_{\text{in}} = 4$. This construction equalizes local branch lengths (and therefore local branch resistances) for edges emanating from a node.

Let $r$ denote channel radius and $S$ denote nozzle spacing. Define geometry-dependent connector-volume constants

$$C_2 = \pi r^2 S, \qquad C_3 = \pi r^2 S\sqrt{3}, \qquad C_4 = 2\sqrt{2}\,\pi r^2 S,$$

#### 3 Combinatorial counts of mixers and connectors

We define the *combinatorial counts* of a hierarchical mixer at refinement depth $\mathcal{L}$ as:

$$N_{\text{mix},n_{\text{in}}}(\mathcal{L}) \quad \text{(number of mixer bodies)}, \qquad N_{\text{con},n_{\text{in}}}(\mathcal{L}) \quad \text{(number of connectors)}.$$

Let $a_{n_{\text{in}}}(\ell) := |V_\ell|$ denote the number of nodes in layer $\ell$:

$$a_2(\ell) = \ell + 1, \qquad a_3(\ell) = \frac{(\ell+1)(\ell+2)}{2}, \qquad a_4(\ell) = (\ell+1)^2.$$

##### 3.1 Degenerate initial layers

Layer $\ell = 1$ consists of inlets and layer $\ell = 2$ consists of single-parent junctions; neither layer contains a mixing element. Therefore, mixer bodies begin at $\ell = 3$:

$$N_{\text{mix},n_{\text{in}}}(\mathcal{L}) = \sum_{\ell=1}^{\mathcal{L}} a_{n_{\text{in}}}(\ell) - \sum_{\ell=1}^{2} a_{n_{\text{in}}}(\ell).$$

Explicitly,

$$\sum_{\ell=1}^{2} a_2(\ell) = 2 + 3 = 5, \qquad \sum_{\ell=1}^{2} a_3(\ell) = 3 + 6 = 9, \qquad \sum_{\ell=1}^{2} a_4(\ell) = 4 + 9 = 13.$$

##### 3.2 Mixer counts

For $\mathcal{L} \geq 2$,

$$N_{\text{mix},2}(\mathcal{L}) = \sum_{\ell=1}^{\mathcal{L}} (\ell+1) - 5 = \frac{\mathcal{L}(\mathcal{L}+3)}{2} - 5,$$

$$N_{\text{mix},3}(\mathcal{L}) = \sum_{\ell=1}^{\mathcal{L}} \frac{(\ell+1)(\ell+2)}{2} - 9 = \frac{\mathcal{L}(\mathcal{L}^2+6\mathcal{L}+11)}{6} - 9,$$

$$N_{\text{mix},4}(\mathcal{L}) = \sum_{\ell=1}^{\mathcal{L}} (\ell+1)^2 - 13 = \frac{(\mathcal{L}+1)(\mathcal{L}+2)(2\mathcal{L}+3)}{6} - 14.$$

##### 3.3 Connector counts

Connectors link adjacent layers and are counted between $\ell$ and $\ell+1$ for $\ell = 1, \ldots, \mathcal{L}-1$. With the present counting convention, the total number of connector *segments* is proportional to the number of nodes in nonterminal layers; we encode this as

$$N_{\text{con},n_{\text{in}}}(\mathcal{L}) = \sum_{\ell=1}^{\mathcal{L}-1} a_{n_{\text{in}}}(\ell) - 1,$$

where the subtraction removes the degenerate inlet connector level in the count convention used for volume accounting. Explicitly,

$$N_{\text{con},2}(\mathcal{L}) = \sum_{\ell=1}^{\mathcal{L}-1} (\ell+1) - 1 = \frac{(\mathcal{L}-1)(\mathcal{L}+2)}{2} - 1,$$

$$N_{\text{con},3}(\mathcal{L}) = \sum_{\ell=1}^{\mathcal{L}-1} \frac{(\ell+1)(\ell+2)}{2} - 1 = \frac{(\mathcal{L}-1)(\mathcal{L}^2+4\mathcal{L}+6)}{6} - 1,$$

$$N_{\text{con},4}(\mathcal{L}) = \sum_{\ell=1}^{\mathcal{L}-1} (\ell+1)^2 - 1 = \frac{\mathcal{L}(\mathcal{L}+1)(2\mathcal{L}+1)}{6} - 2.$$

##### 3.4 Outlet counts and inversion

The outlet count equals the number of nodes in the terminal layer:

$$k = |V_{\mathcal{L}}|.$$

For the three families,

$$k = \begin{cases} \mathcal{L}+1, & n_{\text{in}} = 2, \\ \dfrac{(\mathcal{L}+1)(\mathcal{L}+2)}{2}, & n_{\text{in}} = 3, \\ (\mathcal{L}+1)^2, & n_{\text{in}} = 4. \end{cases}$$

The refinement depth $\mathcal{L}$ can be written as a function of $k$:

$$\mathcal{L}(k) = \begin{cases} k - 1, & n_{\text{in}} = 2, \\ \dfrac{\sqrt{8k+1} - 3}{2}, & n_{\text{in}} = 3, \\ \sqrt{k} - 1, & n_{\text{in}} = 4. \end{cases}$$

##### 3.4.1 Derivation of level–outlet relations.

**Two-input architecture ($n_{\text{in}} = 2$).** The outlet–level relation is

$$k = \mathcal{L} + 1,$$

which immediately yields

$$\mathcal{L}(k) = k - 1.$$

**Three-input architecture ($n_{\text{in}} = 3$).** The outlet count satisfies

$$k = \frac{(\mathcal{L}+1)(\mathcal{L}+2)}{2},$$
$$2k = \mathcal{L}^2 + 3\mathcal{L} + 2.$$

Rearranging gives the quadratic equation

$$\mathcal{L}^2 + 3\mathcal{L} + (2 - 2k) = 0.$$

Solving yields

$$\mathcal{L} = \frac{-3 \pm \sqrt{8k+1}}{2}.$$

Since $\mathcal{L} \geq 0$, the physically admissible solution is

$$\mathcal{L}(k) = \frac{\sqrt{8k+1} - 3}{2}.$$

**Four-input architecture ($n_{\text{in}} = 4$).** The outlet–level relation is

$$k = (\mathcal{L} + 1)^2,$$

which gives

$$\mathcal{L}(k) = \sqrt{k} - 1.$$

##### 3.5 Dead-volume efficiency: closed forms in terms of $k$

Let $V_{\text{mix}}$ denote the internal volume of one mixer body and let $C_2, C_3, C_4$ denote the effective connector-volume constants (units of volume per connector segment). For a given refinement depth $\mathcal{L}$, define

$$V_i(\mathcal{L}) = V_{\text{mix}}\, M_i(\mathcal{L}) + C_i\, C_i^{(\#)}(\mathcal{L}), \qquad i \in \{2, 3, 4\},$$

where $M_i(\mathcal{L})$ is the number of mixers and $C_i^{(\#)}(\mathcal{L})$ is the number of connectors (counts). The dead-volume-per-outlet efficiency is

$$E_i(k) = \frac{V_i(k)}{k}, \qquad V_i(k) := V_i\big(\mathcal{L}(k)\big).$$

Using the combinatorial counts in the main text (mixers and connectors as functions of $\mathcal{L}$), substitution of $\mathcal{L}(k)$ yields the following closed forms.

###### 3.5.1 Two-input family ($n_{\text{in}} = 2$).

With $\mathcal{L} = k - 1$,

$$E_2(k) = \frac{(k^2 + k - 12)\, V_{\text{mix}} + (k^2 - k - 4)\, C_2}{2k}, \qquad V_2(k) = k\, E_2(k).$$

###### 3.5.2 Three-input family ($n_{\text{in}} = 3$).

With $\mathcal{L} = (\sqrt{8k+1} - 3)/2$,

$$E_3(k) = \frac{\left(\frac{k}{6}\sqrt{8k+1} + \frac{k}{2} - 10\right) V_{\text{mix}} + \left(\frac{k}{6}\sqrt{8k+1} - \frac{k}{2} - 2\right) C_3}{k}, \qquad V_3(k) = k\, E_3(k).$$

###### 3.5.3 Four-input family ($n_{\text{in}} = 4$).

With $\mathcal{L} = \sqrt{k} - 1$,

$$E_4(k) = \frac{\left(\frac{\sqrt{k}(\sqrt{k}+1)(2\sqrt{k}+1)}{6} - 14\right) V_{\text{mix}} + \left(\frac{(\sqrt{k}-1)\sqrt{k}(2\sqrt{k}-1)}{6} - 2\right) C_4}{k}, \qquad V_4(k) = k\, E_4(k).$$

###### 3.5.4 Domain of validity.

The efficiency expressions $E_i(k)$ are defined only for outlet counts $k$ that are realizable by the corresponding mixer family. Specifically:

- **Two-input architecture ($i = 2$).** Valid outlet counts satisfy

$$k = \mathcal{L} + 1, \qquad \mathcal{L} \in \mathbb{Z}_{\geq 1}.$$

- **Three-input architecture ($i = 3$).** Valid outlet counts satisfy

$$k = \frac{(\mathcal{L}+1)(\mathcal{L}+2)}{2}, \qquad \mathcal{L} \in \mathbb{Z}_{\geq 1}.$$

- **Four-input architecture ($i = 4$).** Valid outlet counts satisfy

$$k = (\mathcal{L}+1)^2, \qquad \mathcal{L} \in \mathbb{Z}_{\geq 1}.$$

Outside these discrete sequences, the expressions for $E_i(k)$ should be interpreted as formal continuations rather than physically realizable architectures.

#### 3.6 Asymptotic Scaling of Dead-Volume Efficiency

We analyze the asymptotic behavior of the dead-volume-per-outlet efficiency

$$E_i(k) = \frac{V_i(k)}{k},$$

for the $i$-way architectures as the number of outlets $k \to \infty$ along the discrete sequences admissible for each family.

The total dead volume is given by

$$V_i(k) = V_{\text{mix}}\, M_i\big(\mathcal{L}(k)\big) + C_i\, C_i^{(\#)}\big(\mathcal{L}(k)\big),$$

where $\mathcal{L}(k)$ denotes the refinement depth required to realize $k$ outlets, $M_i(\mathcal{L})$ is the number of mixer bodies, and $C_i^{(\#)}(\mathcal{L})$ is the number of connectors.

##### 3.6.1 Two-input architecture ($i = 2$).

For the binary strip, the outlet–level relation is $\mathcal{L} = k - 1$. The mixer and connector counts scale as

$$M_2(\mathcal{L}) = \frac{\mathcal{L}^2}{2} + \mathcal{O}(\mathcal{L}), \qquad C_2^{(\#)}(\mathcal{L}) = \frac{\mathcal{L}^2}{2} + \mathcal{O}(\mathcal{L}).$$

Substituting $\mathcal{L} \sim k$ yields

$$V_2(k) = \frac{(V_{\text{mix}} + C_2)}{2}\, k^2 + \mathcal{O}(k),$$

and therefore

$$E_2(k) = \frac{V_2(k)}{k} = \frac{(V_{\text{mix}} + C_2)}{2}\, k + \mathcal{O}(1).$$

Thus, the dead-volume-per-outlet efficiency of the two-input architecture grows linearly with $k$.

##### 3.6.2 Three-input architecture ($i = 3$).

For the three-input family,

$$\mathcal{L}(k) = \frac{\sqrt{8k+1} - 3}{2} \sim \sqrt{2k} \quad \text{as } k \to \infty.$$

The combinatorial counts satisfy

$$M_3(\mathcal{L}) = \frac{\mathcal{L}^3}{6} + \mathcal{O}(\mathcal{L}^2), \qquad C_3^{(\#)}(\mathcal{L}) = \frac{\mathcal{L}^3}{6} + \mathcal{O}(\mathcal{L}^2).$$

Substituting $\mathcal{L} \sim \sqrt{2k}$ gives

$$V_3(k) = \frac{(V_{\text{mix}} + C_3)}{6} (2k)^{3/2} + \mathcal{O}(k),$$

and hence

$$E_3(k) = \frac{V_3(k)}{k} = \frac{\sqrt{2}}{3} (V_{\text{mix}} + C_3) \sqrt{k} + \mathcal{O}(1).$$

##### 3.6.3 Four-input architecture ($i = 4$).

For the four-input family, the outlet–level relation is

$$\mathcal{L}(k) = \sqrt{k} - 1 \sim \sqrt{k}.$$

The leading-order combinatorial counts are

$$M_4(\mathcal{L}) = \frac{\mathcal{L}^3}{3} + \mathcal{O}(\mathcal{L}^2), \qquad C_4^{(\#)}(\mathcal{L}) = \frac{\mathcal{L}^3}{3} + \mathcal{O}(\mathcal{L}^2).$$

Substitution yields

$$V_4(k) = \frac{(V_{\text{mix}} + C_4)}{3} k^{3/2} + \mathcal{O}(k),$$

and therefore

$$E_4(k) = \frac{V_4(k)}{k} = \frac{1}{3} (V_{\text{mix}} + C_4) \sqrt{k} + \mathcal{O}(1).$$

##### 3.6.4 Summary of asymptotic behavior.

Collecting the leading-order terms,

$$E_2(k) = \Theta(k), \qquad E_3(k) = \Theta(\sqrt{k}), \qquad E_4(k) = \Theta(\sqrt{k}).$$

The two-input architecture exhibits linear growth in dead volume per outlet, whereas the three- and four-input architectures exhibit sublinear $\sqrt{k}$ growth.

##### 3.6.5 Interpretation.

The difference in scaling arises from the dimensionality of the refinement: the two-input architecture refines along a one-dimensional strip, while the three- and four-input architectures refine over two-dimensional planar lattices. As a consequence, the number of internal elements required to realize $k$

outlets grows quadratically for the two-input family but only as $k^{3/2}$ for the higher-valence families, leading to improved asymptotic efficiency.

#### 3.7 Asymptotic Crossover Between Three- and Four-Input Architectures

Although the three- and four-input architectures exhibit the same asymptotic scaling order,

$$E_3(k),\, E_4(k) = \Theta(\sqrt{k}),$$

their leading-order prefactors differ. As a result, the relative ordering of their dead-volume efficiencies depends on the outlet count $k$.

Expanding the closed-form expressions for $E_3(k)$ and $E_4(k)$ to first sub-leading order yields

$$E_3(k) = \frac{\sqrt{2}}{3}\,(V_{\text{mix}} + C_3)\,\sqrt{k} + \frac{1}{2}\,(V_{\text{mix}} - C_3) + O(1),$$

$$E_4(k) = \frac{1}{3}\,(V_{\text{mix}} + C_4)\,\sqrt{k} + \frac{1}{2}\,(V_{\text{mix}} - C_4) + O(1).$$

Setting $E_3(k_c) = E_4(k_c)$ and matching both the $\sqrt{k}$ and $\mathcal{O}(1)$ contributions gives

$$\left[\tfrac{\sqrt{2}}{3}(V_{\text{mix}} + C_3) - \tfrac{1}{3}(V_{\text{mix}} + C_4)\right]\sqrt{k_c} = \tfrac{1}{2}(C_3 - C_4),$$

which solves to

$$k_c = \left[\frac{\tfrac{3}{2}(C_3 - C_4)}{\sqrt{2}(V_{\text{mix}} + C_3) - (V_{\text{mix}} + C_4)}\right]^2.$$

When $\sqrt{2}(V_{\text{mix}} + C_3) < V_{\text{mix}} + C_4$ holds, the three-input architecture exhibits lower dead volume per outlet for $k > k_c$, while the four-input architecture remains more efficient for $k < k_c$. In the opposite regime, in which the mixer volume $V_{\text{mix}}$ dominates the connector contributions $C_3, C_4$, the inequality $\sqrt{2}(V_{\text{mix}} + C_3) > V_{\text{mix}} + C_4$ holds and no physical crossover exists: the four-input architecture is more efficient for all $k$.

##### 3.7.1 Parameter dependence.

The crossover is governed entirely by geometric constants and mixer volume. Since $V_{\text{mix}}$ is fixed while $C_i \propto S$ scales linearly with nozzle spacing $S$, increasing $S$ amplifies the connector contribution relative to the mixer contribution. Consequently, larger nozzle spacings shift the crossover $k_c$ toward smaller outlet counts.

#### 3.8 Geometric feasibility to $n_{\text{in}} \in \{2, 3, 4\}$ families

The constructions above impose the following geometric requirements: (i) the connector primitive is a regular polygonal cell, (ii) the cell tiles the plane, and (iii) the refinement places nodes at cell centers with equal local branch lengths to adjacent vertices. Under these constraints the admissible planar families are based on the line segment ($n_{\text{in}} = 2$), the equilateral triangle ($n_{\text{in}} = 3$), and the square ($n_{\text{in}} = 4$). While a regular hexagon tiles the plane, it cannot recursively subdivided into self-similar hexagons.

Higher valence designs are possible but require the relaxation of one of these constraints.

#### 3.9 Equal-Flow-Per-Layer Flow Model

We assign a flow field to the directed acyclic graph (DAG)

$$G = (V, E),$$

where $V = \{1, \ldots, N\}$ denotes the node set and $E = \{1, \ldots, |E|\}$ denotes the edge set. Each edge $e \in E$ is an ordered pair $(u \to v)$.

The node set is partitioned as

$$I \subset V \quad \text{(inlets)}, \qquad O \subset V \quad \text{(outlets)}, \qquad V_{\text{int}} = V \setminus (I \cup O) \quad \text{(internal nodes)}.$$

A level function $\ell : V \to \mathbb{Z}$ is defined by a breadth-first traversal from the inlets:

$$\ell(i) = 1 \quad \text{for } i \in I, \qquad \ell(v) = \ell(u) + 1 \quad \text{for } (u \to v) \in E.$$

By construction, all outlets lie at level $\ell = n_{\text{Levels}}$.

We prescribe a target inflow $Q_{\text{out}} > 0$ at each outlet. The equal-flow-per-layer model seeks a nonnegative edge-flow vector $Q \in \mathbb{R}^{|E|}$ that:

1. delivers identical total inflow to every outlet,
2. conserves mass at all internal nodes,
3. enforces equal throughput for all nodes within a given internal level, and
4. treats all inlets symmetrically.

Among all such flows, the model selects the one with minimal squared magnitude.

##### 3.9.1 Decision variables.

Each edge $e \in E$ is assigned a flow variable $Q_e \geq 0$, collected as $Q \in \mathbb{R}^{|E|}$. To enforce equal throughput within each internal level, we introduce an auxiliary scalar $T_L$ for each internal level $L$ and define

$$\mathcal{L}_{\text{int}} = \{\, L \mid \exists\, v \in V_{\text{int}} \text{ with } \ell(v) = L \,\}, \qquad n_{\text{lev}} = |\mathcal{L}_{\text{int}}|.$$

The full decision vector is

$$x = \begin{bmatrix} Q \\ T \end{bmatrix} \in \mathbb{R}^{|E| + n_{\text{lev}}}.$$

##### 3.9.2 Outlet constraints.

Each outlet $o \in O$ receives the prescribed inflow:

$$\sum_{e=(u\to o)\in E} Q_e = Q_{\text{out}}. \tag{1}$$

##### 3.9.3 Mass conservation at internal nodes.

For each internal node $v \in V_{\text{int}}$,

$$\sum_{e=(u\to v)\in E} Q_e - \sum_{e=(v\to w)\in E} Q_e = 0. \tag{2}$$

##### 3.9.4 Equal throughput per internal level.

For any internal level $L \in \mathcal{L}_{\text{int}}$, each node $v$ with $\ell(v) = L$ has identical total outflow $T_L$:

$$\sum_{e=(v\to w)\in E} Q_e = T_L. \tag{3}$$

##### 3.9.5 Equal inlet flows.

Let $I = \{i_1, \ldots, i_{n_I}\}$ denote the inlet set. Symmetry is enforced by requiring equal total outflow from all inlets. Taking $i_1$ as a reference, for $k = 2, \ldots, n_I$,

$$\sum_{e=(i_k\to w)\in E} Q_e - \sum_{e=(i_1\to w)\in E} Q_e = 0. \tag{4}$$

##### 3.9.6 Nonnegativity of edge flows.

Backflow is not permitted:

$$Q_e \geq 0 \qquad \forall e \in E. \tag{5}$$

##### 3.9.7 Optimization problem.

The equal-flow-per-layer field is obtained by solving

$$\begin{aligned} &\min_{Q,T} \quad \frac{1}{2}\sum_{e\in E} Q_e^2 \\ &\text{subject to} \quad (1), (2), (3), (4), (5). \end{aligned} \tag{6}$$

In matrix form, with $x = [Q^\top, T^\top]^\top$,

$$\begin{aligned} \min_x \quad & \frac{1}{2} x^\top H x \\ \text{subject to} \quad & A_{\text{eq}} x = b_{\text{eq}}, \\ & x \geq \ell_b, \end{aligned} \tag{7}$$

where $H = \operatorname{diag}(1, \ldots, 1, 0, \ldots, 0)$ penalizes only edge flows, and $\ell_b$ enforces nonnegativity on the first $|E|$ components.

###### 3.9.8 Interpretation.

The resulting flow field satisfies the imposed symmetry and conservation constraints while minimizing overall flow magnitude. It serves as a symmetric reference flow associated with the network topology, rather than a detailed hydrodynamic solution with explicit channel resistances. This reference field provides a consistent baseline for defining and comparing advective mixing operators.

###### 3.9.9 Advective Mixing Matrix for the Equal-Flow-Per-Layer Field

Given the optimized flow $Q^{(\text{layer})}$, we construct an advective mixing operator

$$F_{\text{layer}} \in \mathbb{R}^{|\mathcal{O}| \times |\mathcal{I}|}$$

that maps inlet compositions to outlet compositions.

###### 3.9.10 Directed flow graph.

For each edge $e = (u \to v)$, let

$$q_e = Q_e^{(\text{layer})}.$$

Edges with $q_e = 0$ are discarded. The remaining edges define a directed, feedforward graph.

For each node $j$, define the parent set

$$\mathcal{P}(j) = \{\, i \in V \mid \exists\, e = (i \to j) \text{ with } q_e > 0 \,\},$$

with associated incoming fluxes

$$\phi_{ij} = q_e \quad \text{for } e = (i \to j).$$

###### 3.9.11 Topological ordering.

Since the graph is acyclic, a topological ordering $v_1, \ldots, v_N$ exists. Concentrations are propagated in this order from inlets to outlets.

###### 3.9.12 Advective update rule.

For each inlet $i_k \in \mathcal{I}$, define a concentration field $c^{(k)}$ as follows:

1. **Initialization.**

$$c_{i_k}^{(k)} = 1, \qquad c_i^{(k)} = 0 \quad \forall i \in \mathcal{I} \setminus \{i_k\}.$$

2. **Propagation.** For each non-inlet node $j$,

$$c_j^{(k)} = \begin{cases} \dfrac{\sum_{i \in \mathcal{P}(j)} \phi_{ij} c_i^{(k)}}{\sum_{i \in \mathcal{P}(j)} \phi_{ij}}, & \sum_{i \in \mathcal{P}(j)} \phi_{ij} > 0, \\ c_j^{(k)}, & \text{otherwise.} \end{cases}$$

The mixing matrix is defined by

$$F_{\text{layer}}(o, k) = c_o^{(k)}, \qquad o \in \mathcal{O},\ k = 1, \dots, |\mathcal{I}|.$$

#### 3.10 Binary incomplete Mixing Model

In the two-input architecture, the equal-flow-per-layer solution $Q^{(\text{layer})}$ is treated as a fixed reference flow. An incomplete-mixing rule is applied at each internal $2 \to 2$ node, as detailed below.

#### 3.11 Binary Incomplete Mixing Model

Here we expand the mathematical details of **Fig. 2k ii**, with the binary incomplete mixing model.In the two-input (binary strip) architecture, the equal-flow-per-layer solution $Q^{(\text{layer})}$ is taken as a fixed reference flow field. An incomplete-mixing rule is applied at each internal $2 \to 2$ node.

We work on the directed acyclic graph

$$G = (V, E),$$

with inlet set $\mathcal{I} = \{i_1, i_2\}$, outlet set $\mathcal{O} \subset V$, level function $\ell : V \to \{1, \dots, n_{\text{Levels}}\}$, and edge set $E \subset V \times V$, as defined in the binary case of the general construction. The equal-flow-per-layer model yields a nonnegative edge-flow vector

$$Q^{(\text{layer})} \in \mathbb{R}^{|E|}, \qquad Q_e^{(\text{layer})} \geq 0 \ \ \forall e \in E,$$

such that all outlets receive identical inflow $Q_{\text{out}}$ and all internal nodes satisfy mass conservation and per-level throughput constraints.

##### 3.11.1 Edge compositions.

Each edge $e \in E$ is assigned a scalar composition

$$c_e \in [0, 1],$$

representing the fraction of material originating from inlet $i_1$. Inlet node compositions are initialized as

$$c_{i_1} = 1, \qquad c_{i_2} = 0.$$

All other node compositions are determined by upstream propagation.

For any node $v \in V$, define the sets of incoming and outgoing edges

$$\mathcal{E}^{\mathrm{in}}(v) = \{e = (u \to v) \in E\}, \qquad \mathcal{E}^{\mathrm{out}}(v) = \{e = (v \to w) \in E\}.$$

The algorithm proceeds by sweeping levels $L = 1, \ldots, n_{\mathrm{Levels}} - 1$, updating compositions on $\mathcal{E}^{\mathrm{out}}(v)$ for each node $v$ at level $L$ according to the following cases.

##### 3.11.2 Case 1: inlet nodes.

If $\mathcal{E}^{\mathrm{in}}(v) = \emptyset$, the node composition is prescribed externally and copied to all outgoing edges:

$$c_e = c_v, \qquad \forall e \in \mathcal{E}^{\mathrm{out}}(v).$$

##### 3.11.3 Case 2: single-parent nodes.

If $|\mathcal{E}^{\mathrm{in}}(v)| = 1$, no mixing occurs. Let $\mathcal{E}^{\mathrm{in}}(v) = \{e_{\mathrm{par}}\}$. Then

$$c_e = c_{e_{\mathrm{par}}}, \qquad \forall e \in \mathcal{E}^{\mathrm{out}}(v).$$

##### 3.11.4 Case 3: incomplete mixing at a 2 → 2 node.

If

$$|\mathcal{E}^{\mathrm{in}}(v)| = 2, \qquad |\mathcal{E}^{\mathrm{out}}(v)| = 2,$$

the node admits incomplete mixing. Label the incoming edges $\mathcal{E}^{\mathrm{in}}(v) = \{e_L, e_R\}$ by sorting parent-node indices, with

$$q_L = Q^{(\mathrm{layer})}_{e_L}, \qquad q_R = Q^{(\mathrm{layer})}_{e_R},$$

and compositions

$$c_L = c_{e_L}, \qquad c_R = c_{e_R}.$$

The total inflow and perfectly mixed composition at node $v$ are

$$Q_{\mathrm{in}} = q_L + q_R, \qquad c_{\mathrm{mix}} = \frac{q_L c_L + q_R c_R}{Q_{\mathrm{in}}}.$$

The outgoing edges are labeled $\mathcal{E}^{\mathrm{out}}(v) = \{e_{\mathrm{left}}, e_{\mathrm{right}}\}$ by sorting child-node indices. incomplete mixing is parameterized by $\alpha \in [0, 1]$, where $\alpha = 0$ corresponds to no mixing and $\alpha = 1$ to perfect mixing. The child-edge compositions are given by

$$c_{e_{\text{left}}} = \alpha\, c_{\text{mix}} + (1-\alpha)\, c_L, \tag{8}$$

$$c_{e_{\text{right}}} = \alpha\, c_{\text{mix}} + (1-\alpha)\, c_R. \tag{9}$$

##### 3.11.5 Outlet compositions.

After processing all internal levels, each outlet $o \in \mathcal{O}$ receives flow through incoming edges $\mathcal{E}^{\text{in}}(o)$. The outlet composition is defined as the flow-weighted average

$$c_{\text{out}}(o) = \frac{\sum_{e\in\mathcal{E}^{\text{in}}(o)} Q_e^{(\text{layer})}\, c_e}{\sum_{e\in\mathcal{E}^{\text{in}}(o)} Q_e^{(\text{layer})}}, \qquad o \in \mathcal{O}.$$

### 4 Supplementary Video Descriptions

**Supplementary Video 1** – 3D gradient multinozzle embedded printing of trileaflet valves. Here we show pressure based extrusion printing of tri-leaflet valves with a GEM three-way printhead with ink (ETD 2020 0.3 wt%) colored with different parent fluorescent pigments (Yellow, Blue, Pink) from isometric, bottom, and frontal views.

**Supplementary Video 2** – .Hemodynamic testing under pulmonary pressures and flows of 3D-printed valve made with 20k/35k/8-arm PEGDA mix optimized using three-way GEM printhead.

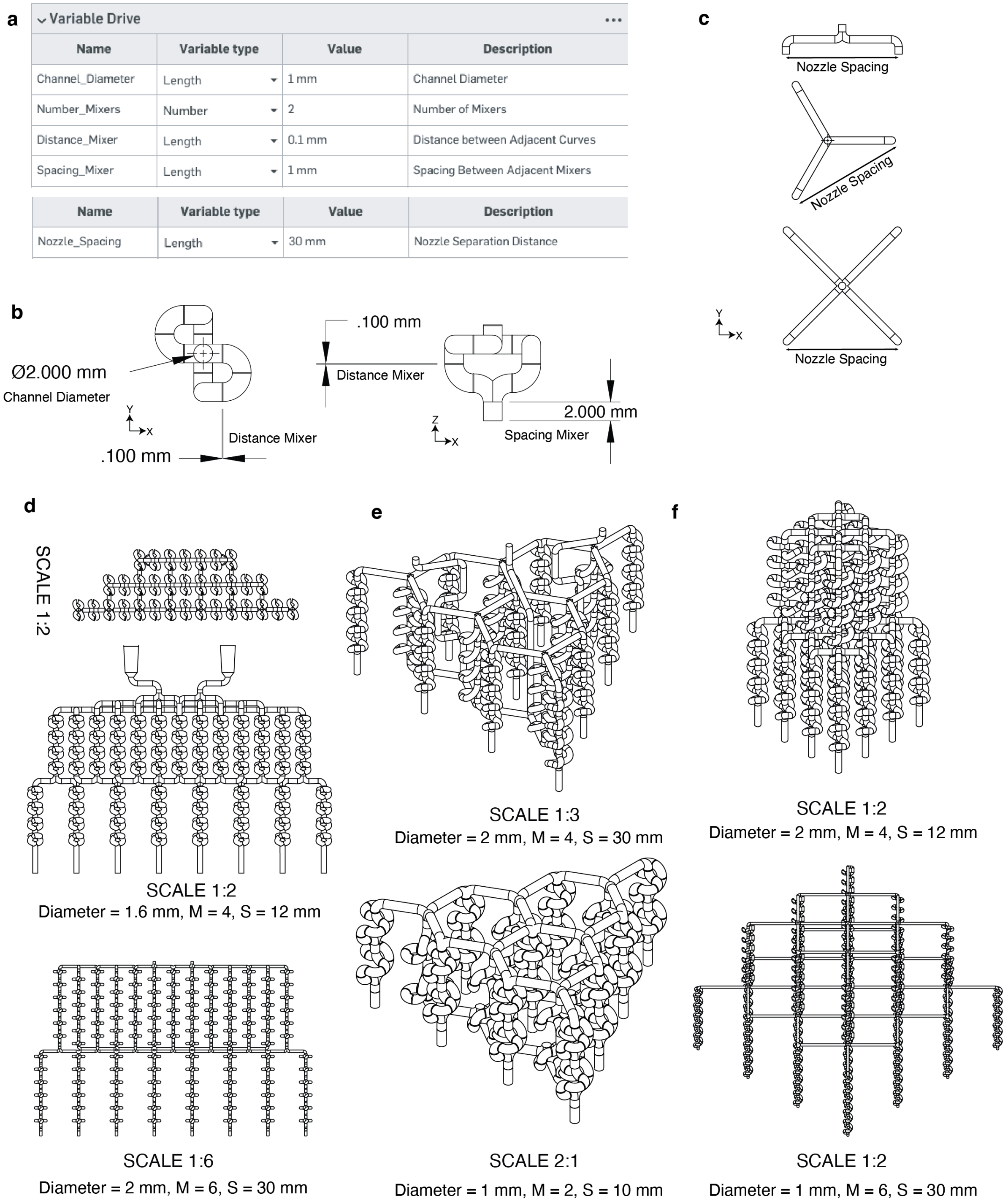


**Fig. 1: Construction of GEM nozzles using variable-driven computer-aided design. a**, Variable tables defining the parametric CAD framework used to generate GEM printhead architectures. Nozzle separation distance, channel diameter, number of mixers ($M$), inter-mixer spacing, and intra-mixer curvature offsets are defined through global design variables. **b**, Dimensioned planar views of a single mixing element showing key geometric parameters. **c**, Schematic representations of outlet nozzle separation distance for two-, three-, and four-way printhead configurations, illustrating how nozzle spacing is parameterized independently of the internal mixing architecture. **d–f**, Example assemblies and isometric and planar views of two-, three-, and four-way GEM printheads generated by hierarchical stacking of mixing elements across multiple layers and length scales, demonstrating scalability of the design framework across channel diameters, number of mixers ($M$), and outlet spacing ($S$).

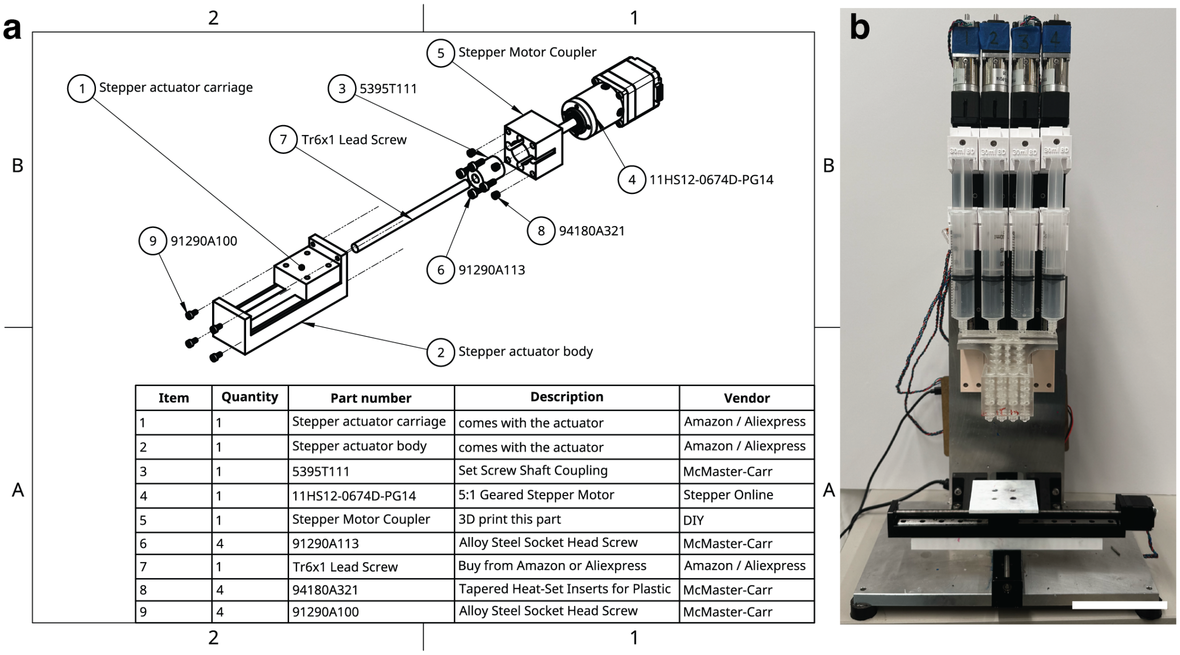


| Item | Quantity | Part number | Description | Vendor |
|---|---|---|---|---|
| 1 | 1 | Stepper actuator carriage | comes with the actuator | Amazon / Aliexpress |
| 2 | 1 | Stepper actuator body | comes with the actuator | Amazon / Aliexpress |
| 3 | 1 | 5395T111 | Set Screw Shaft Coupling | McMaster-Carr |
| 4 | 1 | 11HS12-0674D-PG14 | 5:1 Geared Stepper Motor | Stepper Online |
| 5 | 1 | Stepper Motor Coupler | 3D print this part | DIY |
| 6 | 4 | 91290A113 | Alloy Steel Socket Head Screw | McMaster-Carr |
| 7 | 1 | Tr6x1 Lead Screw | Buy from Amazon or Aliexpress | Amazon / Aliexpress |
| 8 | 4 | 94180A321 | Tapered Heat-Set Inserts for Plastic | McMaster-Carr |
| 9 | 4 | 91290A100 | Alloy Steel Socket Head Screw | McMaster-Carr |

**Fig. 2: Assembly schematic of the high-torque adapter and extrusion printer mounting. a**, Assembly schematic and bill of materials for the high-torque adapter used for extrusion-based printing, including part numbers, vendors, and assembly instructions. **b**, Four-way GEM printhead mounted on the extrusion printer based on the low-cost Printess bioprinter, modified with four high-torque adapters to enable simultaneous extrusion (Scale bar, 10 cm).

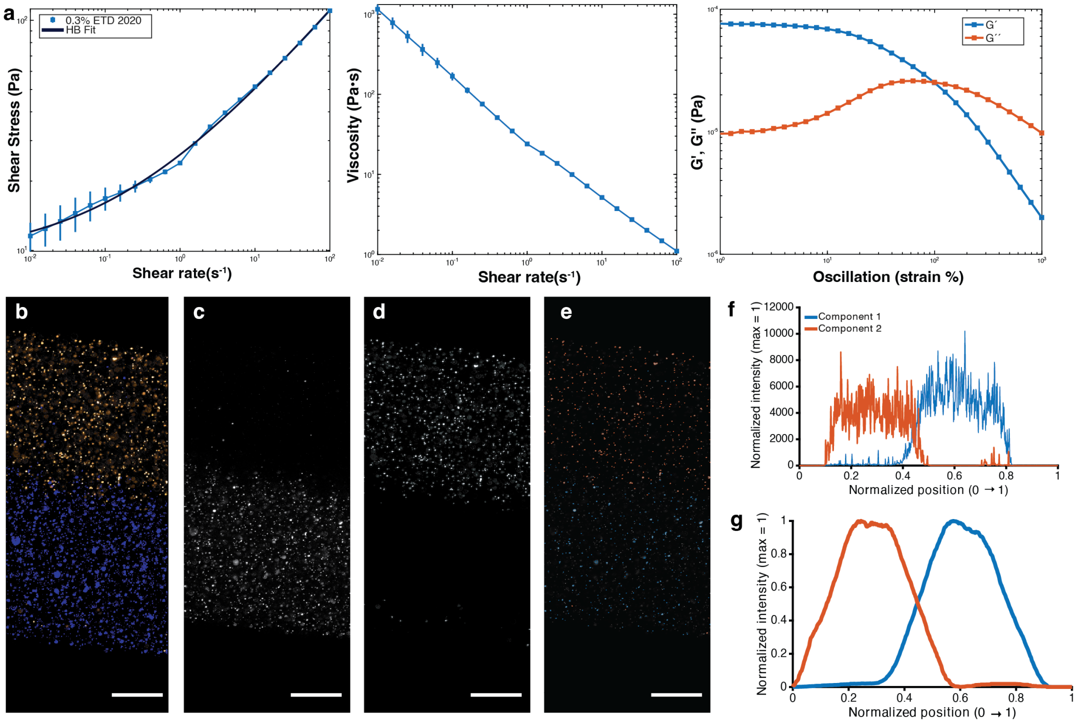


**Fig. 3: Characterization of ink rheology and degree of mixing for Carbopol inks flowing through GEM mixers.** **a**, Rheological characterization of 0.3 wt% ETD 2020 Carbopol ink. Left: shear stress versus shear rate with Herschel–Bulkley (HB) model fit. Middle: viscosity as a function of shear rate, illustrating shear-thinning behavior. Right: oscillatory rheology showing storage ($G'$) and loss ($G''$) moduli as a function of strain amplitude, with the crossover point indicating yield-stress behavior. **b**–**e**, Representative confocal ($10\times$) projections of filaments printed using increasing numbers of stacked mixing elements, containing fluorescent microspheres as tracers to visualize spatial distribution of component streams. Individual channels are thresholded using global Otsu thresholding, and particles are identified using `bwareaopen`, with segmented particle outlines shown (Scale bar = 250 $\mu$m). **f**, Raw diametrical fluorescence intensity profiles extracted across the filament width, plotted as a function of normalized lateral position. **g**, Smoothed and normalized intensity profiles derived from panel **f** using a moving window of 50 pixels.

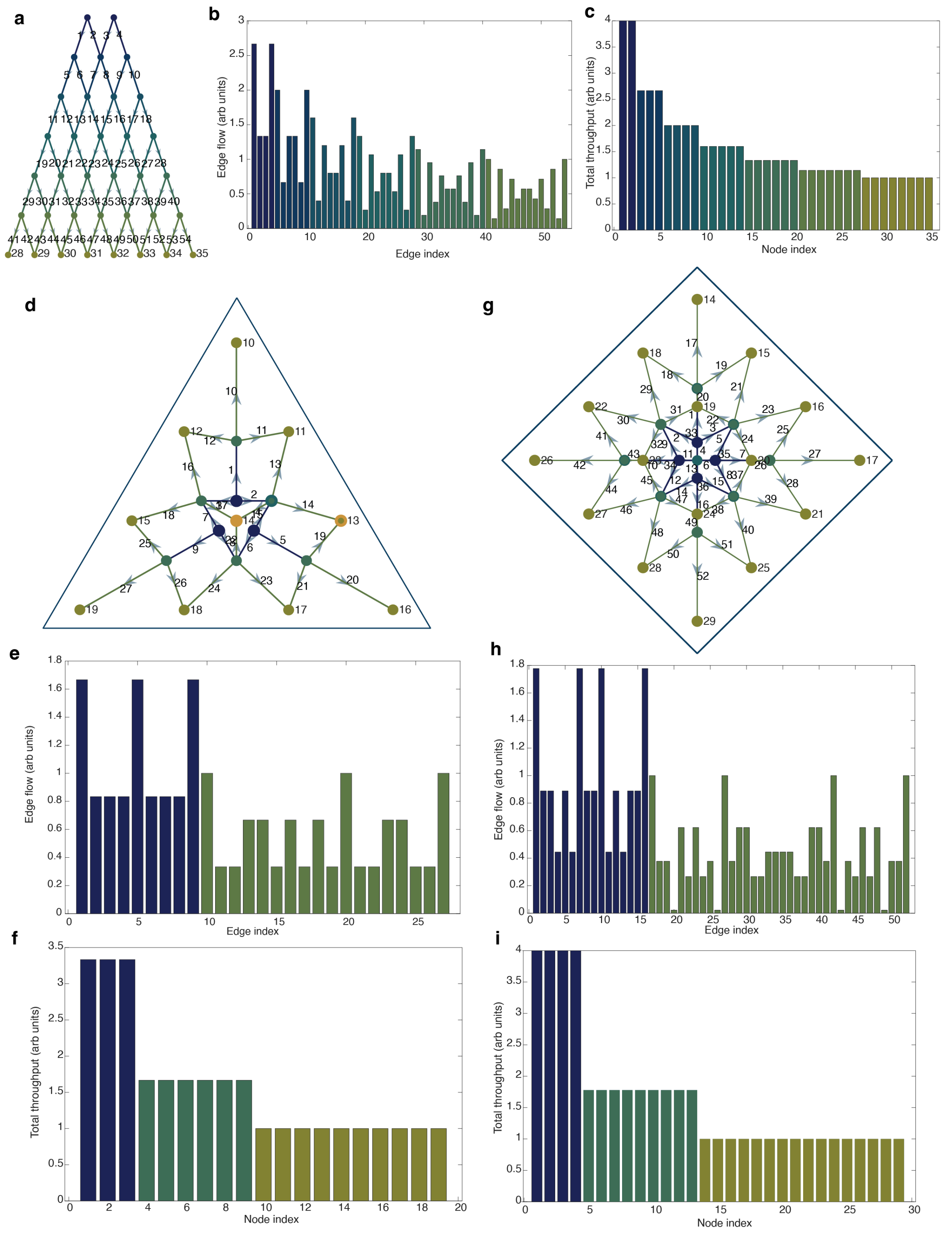


**Fig. 4: Theoretical ink compositions in GEM printheads. a–i**, Directed acyclic graph (DAG) representations of two-way, three-way, and four-way GEM printhead architectures. Edges are weighted by the equal-flow-per-layer solution and colored to indicate layer index, progressing from inlet layers (blue) to outlet layers (light green), illustrating theoretical propagation of ink compositions through the network.

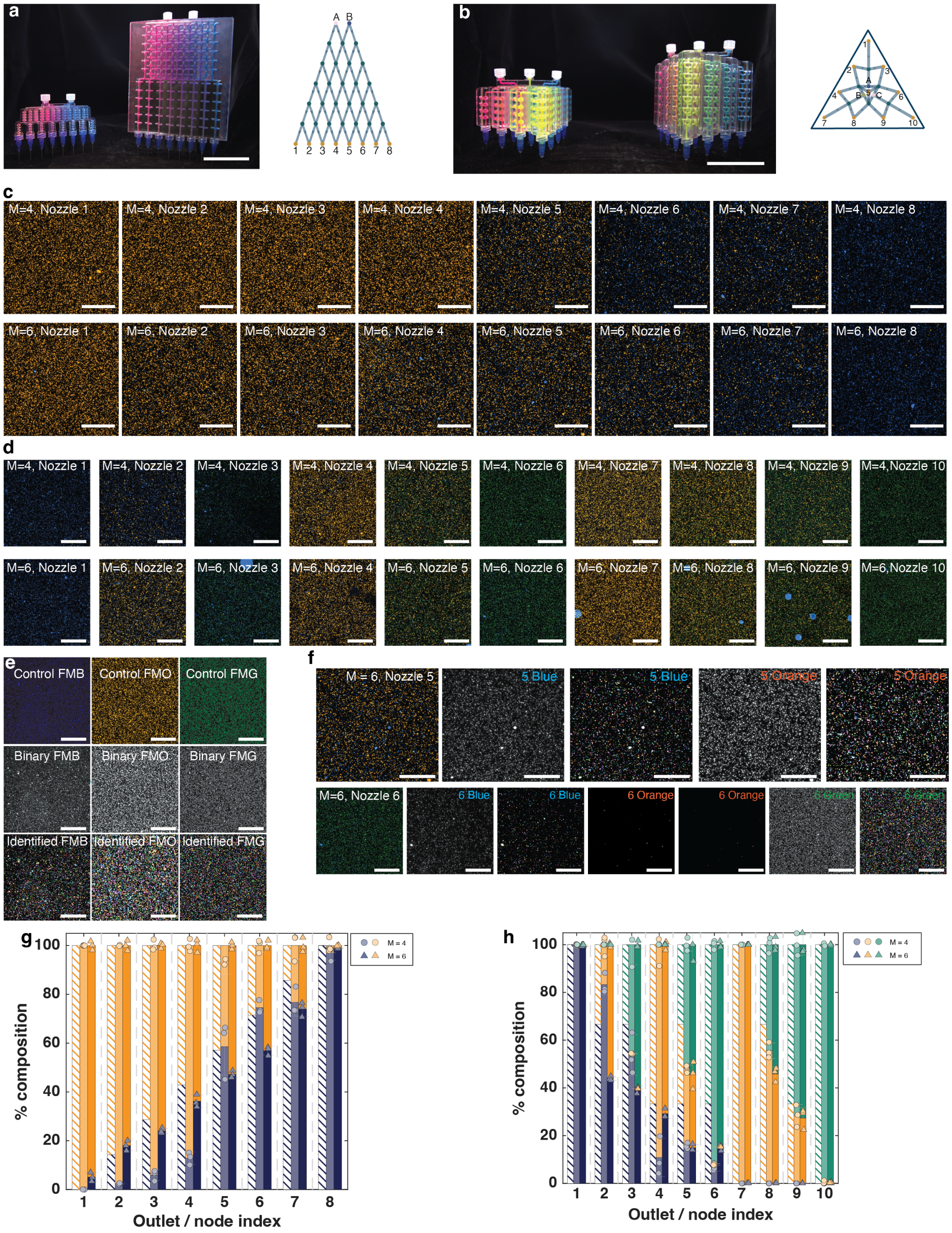

**Fig. 5: Confocal imaging characterization of two- and three-way GEM printheads.** **a–b**, Images of fluorescent pigment–filled two-way (**a**) and three-way (**b**) GEM printheads, shown with $M = 4$ mixers (left) and $M = 6$ mixers (right) (Scale bar, 5 cm). **c–d**, Representative confocal images of fluorescent microspheres collected at each nozzle outlet for both mixer variants and printhead families (Scale bar, 200 $\mu$m). **e**, Parent inks prior to printing, imaged by confocal microscopy, thresholded using Otsu's method, and segmented into individual particles to establish a control baseline particle count for each ink (Scale bar, 200 $\mu$m). **f**, For each nozzle outlet, confocal images were split into their respective spectral channels, thresholded using thresholds determined from the control inks, and segmented to identify particles. Particle overlays are shown. Particle counts for each channel were calculated and normalized using the corresponding control values (Scale bar, 200 $\mu$m). **g–h**, Quantified outlet compositions for two-way and three-way GEM printheads with $M = 4$ and $M = 6$ mixers ($n = 3$), with theoretical predictions shown as striped bars and experimental measurements shown as solid bars.

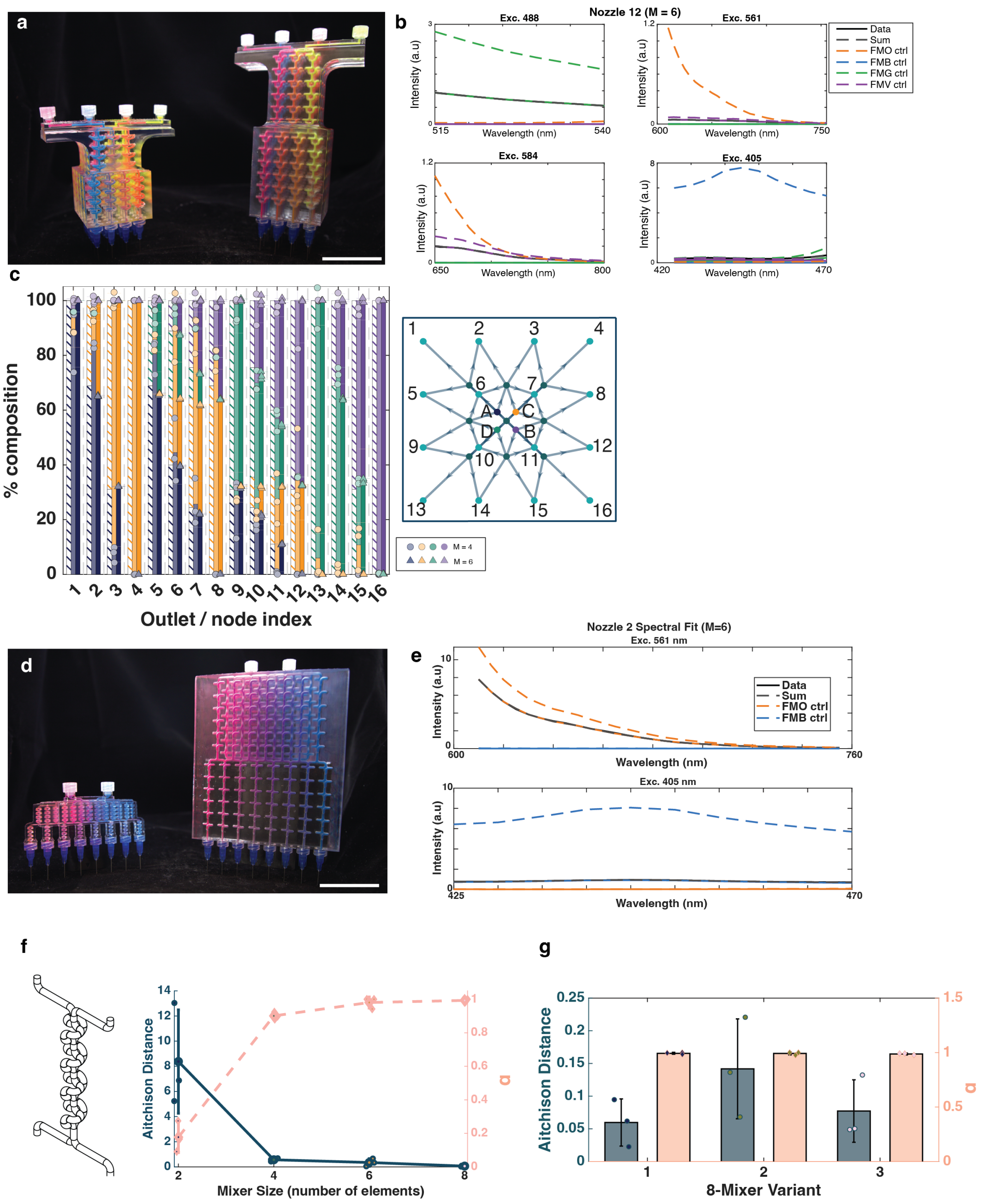


**Fig. 6: Spectral unmixing characterization of four-way GEM printheads and binary incomplete mixing in two-way printheads.** **a**, Images of fluorescent pigment–filled four-way GEM printheads with $M = 4$ mixers (left) and $M = 6$ mixers (right) (Scale bar, 5 cm). **b**, Representative emission spectra acquired under multiple excitation wavelengths, together with the corresponding linear spectral unmixing components and reconstructed signal. **c**, Quantified outlet compositions for four-way GEM printheads with $M = 4$ and $M = 6$ mixers ($n = 3$), with theoretical predictions shown as striped bars. **d–e**, Two-way GEM printheads with $M = 4$ and $M = 6$ mixers analyzed using spectral unmixing to assess binary incomplete mixing (Scale bar, 5 cm). **f**, Isometric view of an isolated two-channel, two-outlet node from a two-way printhead with $M = 4$ mixers. **g**, Quantification of mixing performance as a function of mixer number using Aitchison distance (blue) and the binary imperfect mixing parameter $\alpha$ (pink) ($n = 3$). **h**, Mixing performance metrics (Aitchison distance and $\alpha$) for multiple eight-mixer printhead variants, demonstrating uniformity and reproducibility across independently printed mixers.

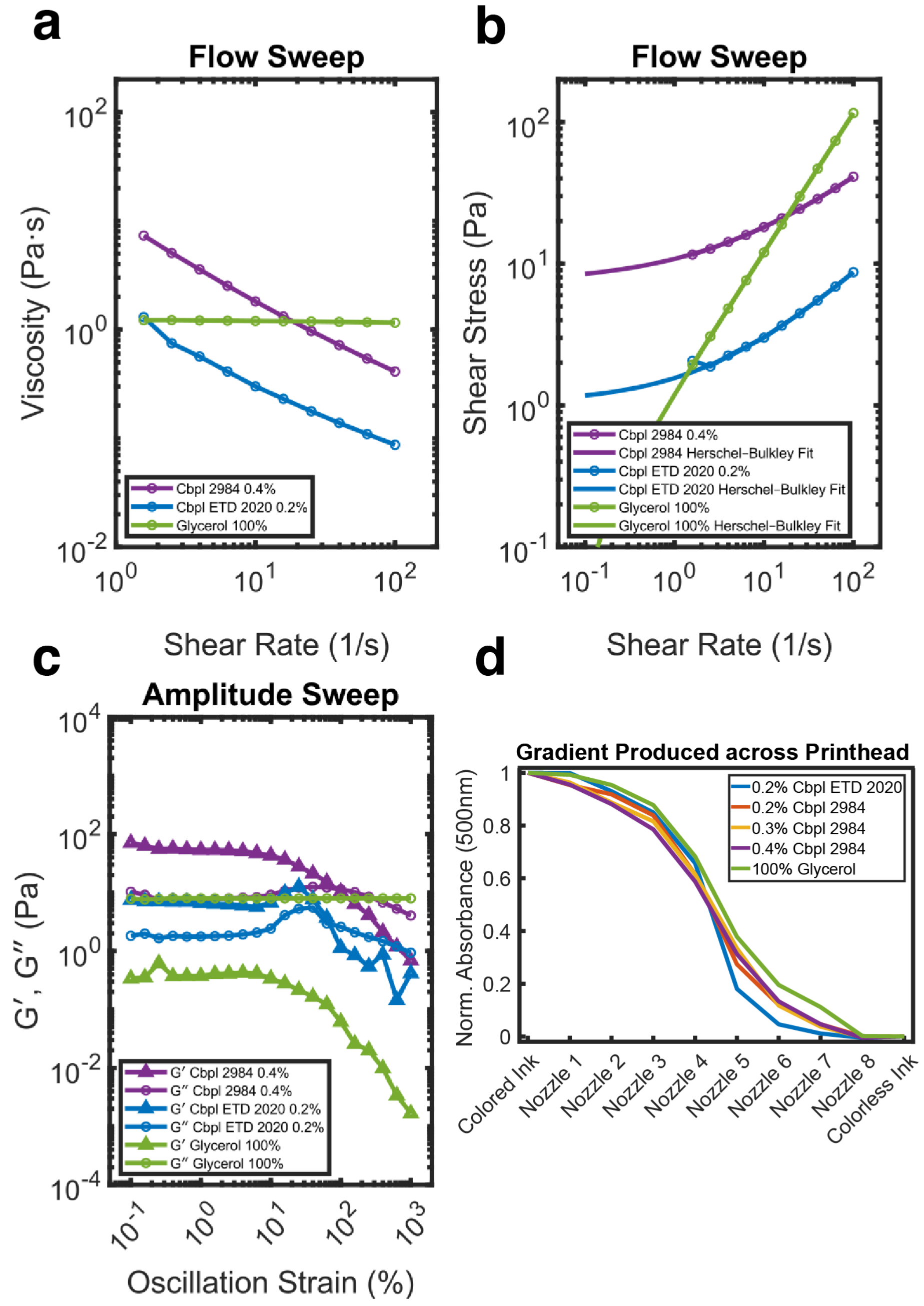


**Fig. 7: Two-way GEM printhead performance for Newtonian, shear-thinning, and yield-stress fluids.** Three inks with distinct rheological behaviors were characterized: two shear-thinning yield-stress fluids (0.4 wt% Carbopol 2984 and 0.2 wt% Carbopol ETD 2020) and one Newtonian fluid (glycerol). **a**, Viscosity as a function of shear rate. **b**, Shear stress versus shear rate with corresponding Herschel–Bulkley model fits. **c**, Oscillatory amplitude sweeps showing storage ($G'$) and loss ($G''$) moduli as a function of strain amplitude. **d**, Outlet concentration distributions for a two-way GEM printhead employing $M = 4$ mixers per node for each ink type. In all cases, identical inks were loaded into both inlets, with fluorescent dye added to one inlet to enable concentration tracking across the outlet array.

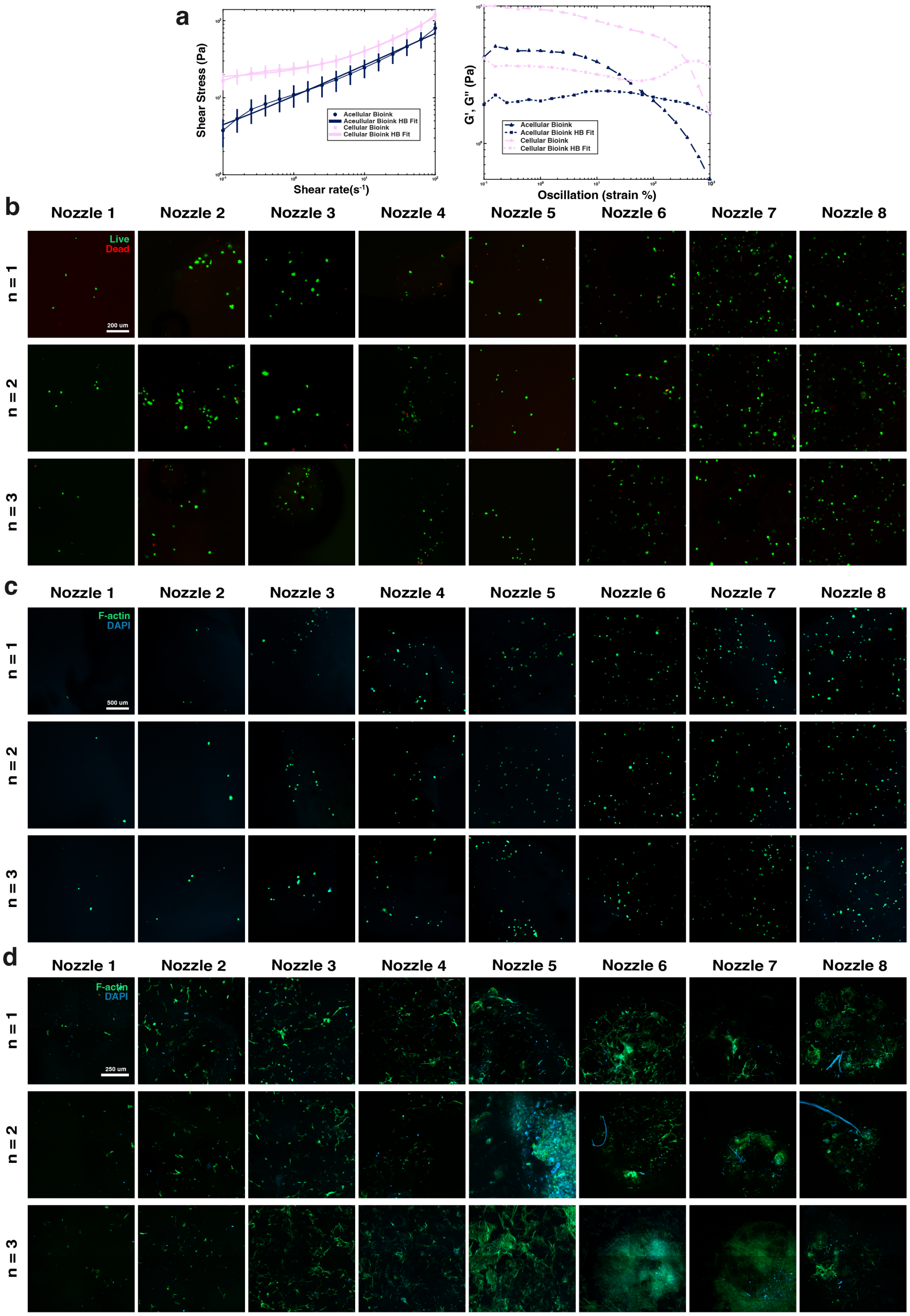


**Fig. 8: Extended bioink rheology and confocal imaging characterization.** **a**, Rheological characterization of acellular and cellular bioinks. Left: shear stress as a function of shear rate. Right: storage ($G'$) and loss ($G''$) moduli measured by oscillatory rheology. **b**–**d**, Confocal microscopy images collected across all eight nozzles and $n = 3$ biological replicates. **b**, Post-print live/dead viability assay. **c**, Post-print F-actin and DAPI staining for cell density characterization. **d**, F-actin and DAPI staining on day 5 to assess scaffold compaction. Scale bars, 200 $\mu$m (**b**), 500 $\mu$m (**c**), and 250 $\mu$m (**d**).

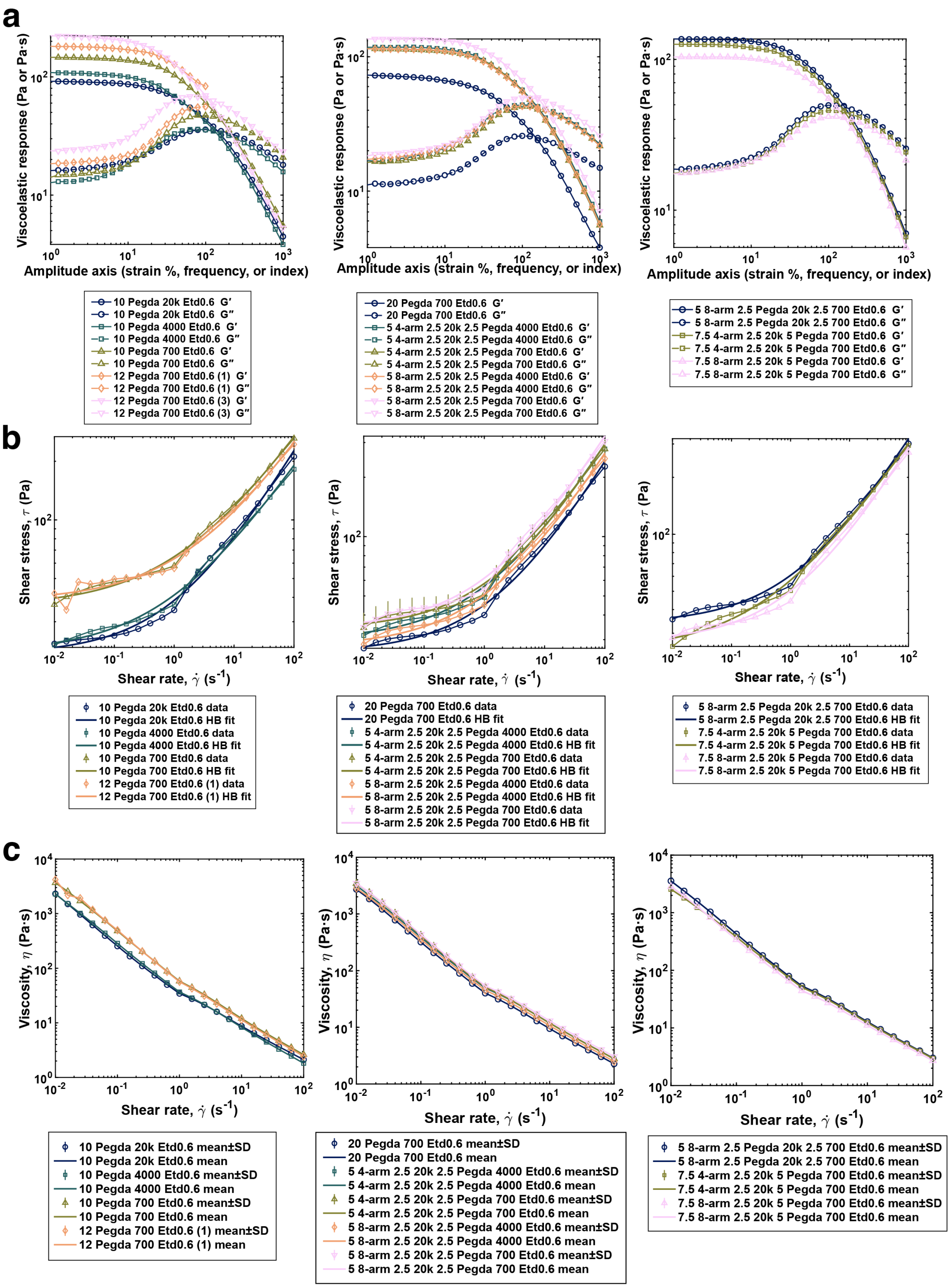


**Fig. 9: Rheological characterization of 10 wt% and 20 wt% PEGDA ink families. a**, Complex viscosity as a function of strain for the different PEGDA formulation families evaluated using the three-way GEM printhead. **b**, Herschel–Bulkley model fits to the rheology data. **c**, Flow sweeps of the different PEGDA-based inks, illustrating shear-thinning behavior

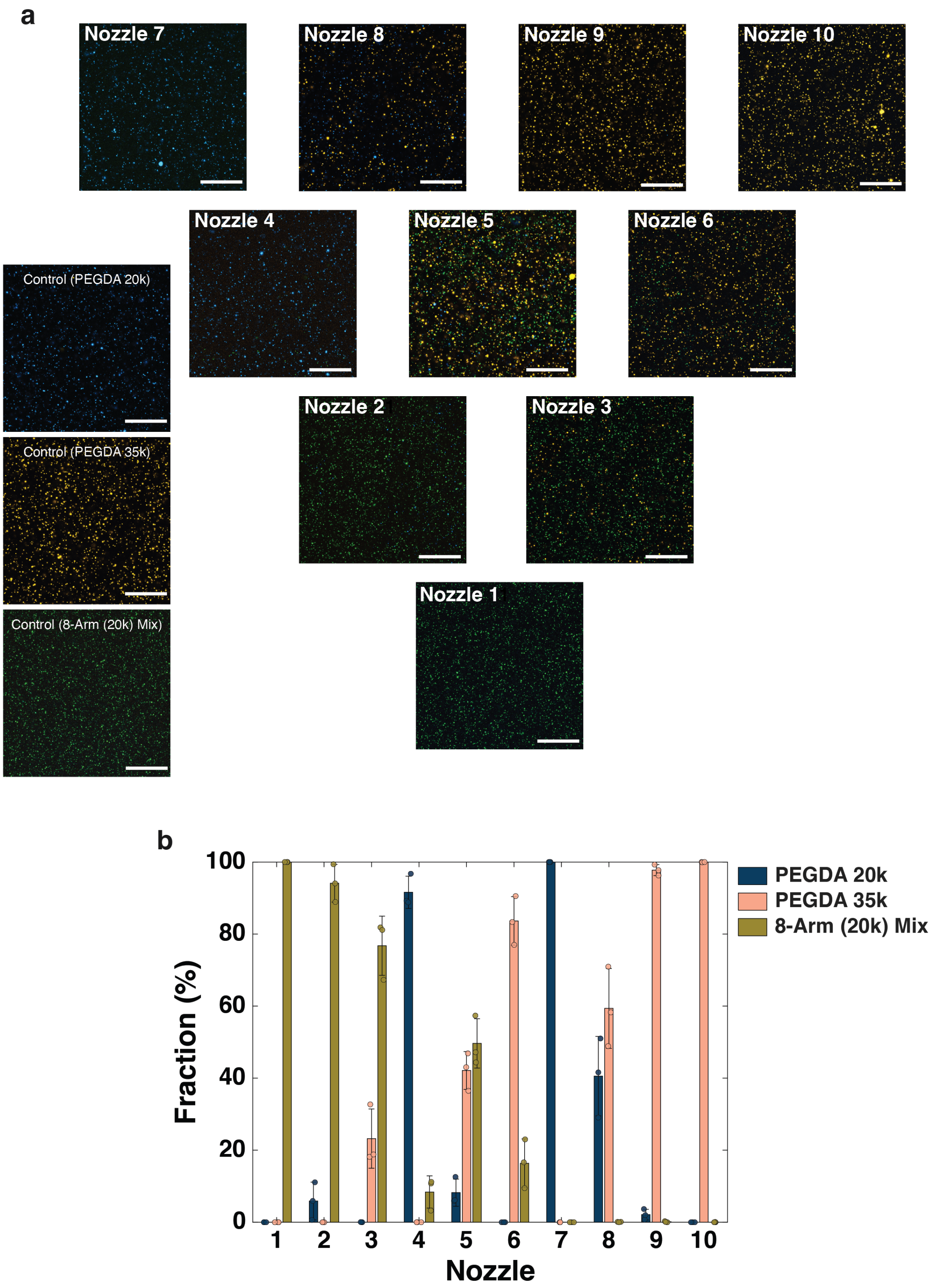


**Fig. 10: Confocal of Output of Valves.** **a**, Representative Confocal images used to identify composition of the input inks (left, 'control') and the outputs from each nozzle. (Scale bar, 200 $\mu$m) **b**, Composition of ink from each nozzle normalized to input inks.

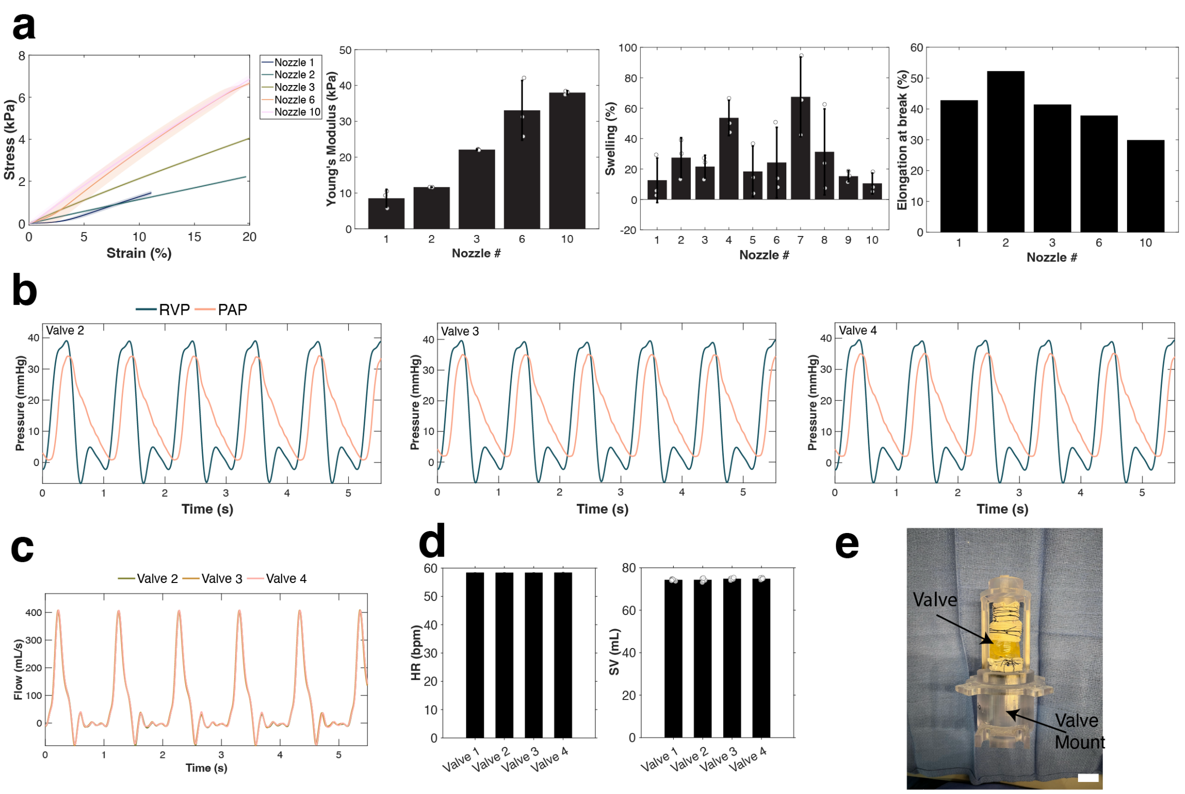


**Fig. 11: Material characterization of 4k/20k/8-arm PEGDA mix and comprehensive hemodynamic valve data from nozzle five of 20k/35k/8-arm PEGDA. a**, Stress–strain uniaxial tensile data ($n = 3$) of 4k/20k/8-arm mix, Young's modulus ($n = 3$), swelling ratio ($n = 3$), and elongation to failure ($n = 1$) (left to right). Only material combinations from nozzles numbers 1, 2, 3, and 10 could be handled and were strong enough to undergo mechanical testing. **b**, Superimposed right ventricular pressure (RVP) and pulmonary arterial pressure (PAP) over six consecutive cycles for valves 2–4 (from nozzle 5 of 20k/35k/8-arm PEGDA). **c**, Flow waveforms over six consecutive cycles for valves 2–4. **d**, Heart rate (HR) and stroke volume (SV) used for hemodynamic testing for all four valves. **e**, Photography of the valve attached to the valve mount prior to hemodynamic testing (Scale bar, 1 cm).
